# Engineered unnatural ubiquitin for optimal detection of deubiquitinating enzymes

**DOI:** 10.1101/2020.01.30.926881

**Authors:** Wioletta Rut, Mikołaj Żmudziński, Scott J. Snipas, Miklos Bekes, Tony T. Huang, Marcin Drag

## Abstract

Deubiquitinating enzymes (DUBs) are responsible for removing ubiquitin (Ub) from its protein conjugates. DUBs have been implicated as attractive therapeutic targets in the treatment of viral diseases, neurodegenerative disorders and cancer. The lack of selective chemical tools for the exploration of these enzymes significantly impairs the determination of their roles in both normal and pathological states. Commercially available fluorogenic substrates are based on the C-terminal Ub motif or contain Ub coupled to a fluorophore (Z-LRGG-AMC, Ub-AMC); therefore, these substrates suffer from lack of selectivity. By using a hybrid combinatorial substrate library (HyCoSuL) and a defined P2 library containing a wide variety of nonproteinogenic amino acids, we established a full substrate specificity profile for two DUBs—MERS PLpro and human UCH-L3. Based on these results, we designed and synthesized Ub-based substrates and activity-based probes (ABPs) containing selected unnatural amino acids located in the C-terminal Ub motif. Biochemical analysis and cell-based experiments confirmed the activity and selectivity of engineered Ub-based substrates and probes. Using this approach, we propose that for any protease that recognizes Ub and Ub-like substrates, a highly active and selective unnatural substrate or probe can be engineered.

## 1. Introduction

Ubiquitin (Ub) conjugation is one of the most important posttranslational modifications and influences the activity, protein-protein interactions, localization and stability of many proteins [1]. Attachment of Ub to protein substrates is a covalent modification mediated by the E1, E2 and E3 enzymes. However, due to the activity of deubiquitinating enzymes (DUBs), this modification is reversible. DUBs hydrolyze the isopeptide bond between the C-terminal glycine of Ub and its substrate or between Ubs in poly-Ub chains with various topologies [2]. The human genome encodes approximately 100 DUBs divided into six subclasses based on sequence similarity and mechanism of action. The majority of DUBs (five of the six subclasses) are cysteine proteases, and only one subclass is represented by a small group of metalloproteases. The largest and most structurally diverse subclass of DUBs are Ub-specific proteases (USPs). These enzymes can hydrolyze a peptide/isopeptide bond between the C-terminus of Ub and its substrate, as well as within the poly-Ub chain. Another subclass of DUBs that belongs to cysteine proteases are the Ub carboxy-terminal hydrolase (UCH) enzymes responsible for the hydrolysis of Ub precursors, leading to the formation of Ub monomers. The other subclasses of cysteine proteases are the ovarian tumor (OTU), Machado-Josephine domain (MJD), and motif-interacting with Ub-containing novel DUB family (MINDY) proteases and one metalloprotease subgroup, known as the JAMM/MPN+ DUBs. Members of the OTU family exhibit unique specificity for cleavage of different poly-Ub chain topologies [3]. Furthermore, enzymes displaying deubiquitinating activity are encoded in coronavirus genomes, such as Middle East respiratory syndrome papain-like protease (MERS PLpro) [4] or severe acute respiratory syndrome papain-like protease (SARS PLpro) [5]. PLpro enzymes are involved in negative regulation of the innate immune response during viral infection [4].

Since DUBs are highly specific toward Ub, the most commonly used DUB assay reagents are based on Ub with a C-terminal fluorogenic group (Ub-ACC [6], Ub-AMC [7], Ub-rhodamine [8]) or an electrophilic warhead (Ub-CHO, Ub-VS, Ub-vinyl methyl ester (VME) [9-12]). Application of these chemical tools in biological studies has led to a better understanding of the mechanisms of action of many DUBs. Ub with a C-terminal aldehyde group has been frequently used in crystallographic studies of DUBs [13]. Development of Ub-based probes by introducing a detection tag on the N-terminus and a reactive group on the C-terminus of Ub enabled labeling of DUBs in cell lysates and their detection via Western blot analysis. Ub-based probes are powerful tools for characterizing the activity of DUBs, identifying new DUBs, evaluating the selectivity of DUB inhibitors, and determining the functional roles of DUBs in normal and pathological states [12, 14]. However, the use of mono-Ub probes precluded studies of DUB activity and linkage specificity. To overcome this limitation, di-Ub probes were developed [15, 16]. Studies with di-Ub probes revealed differences between the S1-S1’ and S1-S2 selectivity of enzymes from the OTU subclass [17], as well as the Lys48 linkage specificity of SARS PLpro [18, 19]. Despite the wide utility of broad-spectrum DUB substrates and probes in biochemical assays, selective chemical tools to study a single DUB (or a narrow subset of DUBs) are needed. Recently, the Ovaa group developed Ub-based probes selective toward USP7 [20] and USP16 [21] by combining structural analysis, modeling and mutational predictions. Selectivity can also be achieved by modifying the Ub C-terminal conserved LRGG motif [22]. Determination of DUB substrate preferences in the P4-P2 positions using the positional scanning substrate combinatorial library (PS-SCL) approach revealed that DUBs can recognize amino acids other than the canonical LRG in these positions. Furthermore, this study shows that a Ub mutant with C-terminal extension, in which glycine in the P2 position was replaced by valine, was cleaved by UCH-L3 20-50-fold less efficiently than its wild-type analog [22]. However, this mutation provided selectivity, as the mutant was not recognized by another tested DUB USP5 (IsoT). In the present study, to further improve Ub mutant selectivity, as well as DUB activity, we decided to utilize unnatural amino acids. A similar approach was successfully applied to design selective substrates and activity-based probes (ABPs) for cysteine [23, 24], serine [25] and threonine proteases [26] as short peptide sequences but never incorporated into whole substrate proteins, such as Ub.

In this work, we developed active and selective chemical tools based on Ub scaffolds for DUB investigation by incorporating unnatural amino acids into the conserved C-terminal Ub motif. We selected two DUBs, namely, UCH-L3 and MERS PLpro, to validate our chemical approach. These enzymes represent potential therapeutic targets in drug design [4, 27, 28]. First, we determined full substrate specificity profiles of the selected DUBs at the P4-P2 positions using defined and hybrid combinatorial libraries of tetrapeptide fluorogenic substrates. Substrate preference analysis allowed us to extract key amino acids that could serve as selectivity-providing motifs in Ub-based substrates and ABPs. The designed tetrapeptide fluorogenic substrates containing unnatural amino acids were selective toward UCH-L3 and MERS PLpro but still not efficiently hydrolyzed. To overcome this obstacle, we synthesized Ub derivatives containing a C-terminal epitope substituted with selected unnatural peptide sequences instead of a canonical LRGG motif. Substrates and covalent probes were obtained by conjugation of the fluorescent label (ACC) or Michael-acceptor warhead (VME), respectively (**Figure 1**). Kinetic analysis revealed that the designed substrates were more efficiently hydrolyzed than the reference “wild-type” substrate Ub-ACC. Moreover, the substrates exhibited high selectivity toward UCH-L3 and MERS PLpro. The selectivity of the Ub-based probe was evaluated using purified recombinant DUBs and cell lysates. The results of these studies revealed that engineered B-U1-VME and B-U2-VME probes detected UCH-L3 in cell lysates with high selectivity. With this *proof of concept*, we propose that this approach can be successfully applied in the design of Ub-based selective chemical tools for other proteases that recognize Ub and Ub-like substrates.

**Figure 1.**
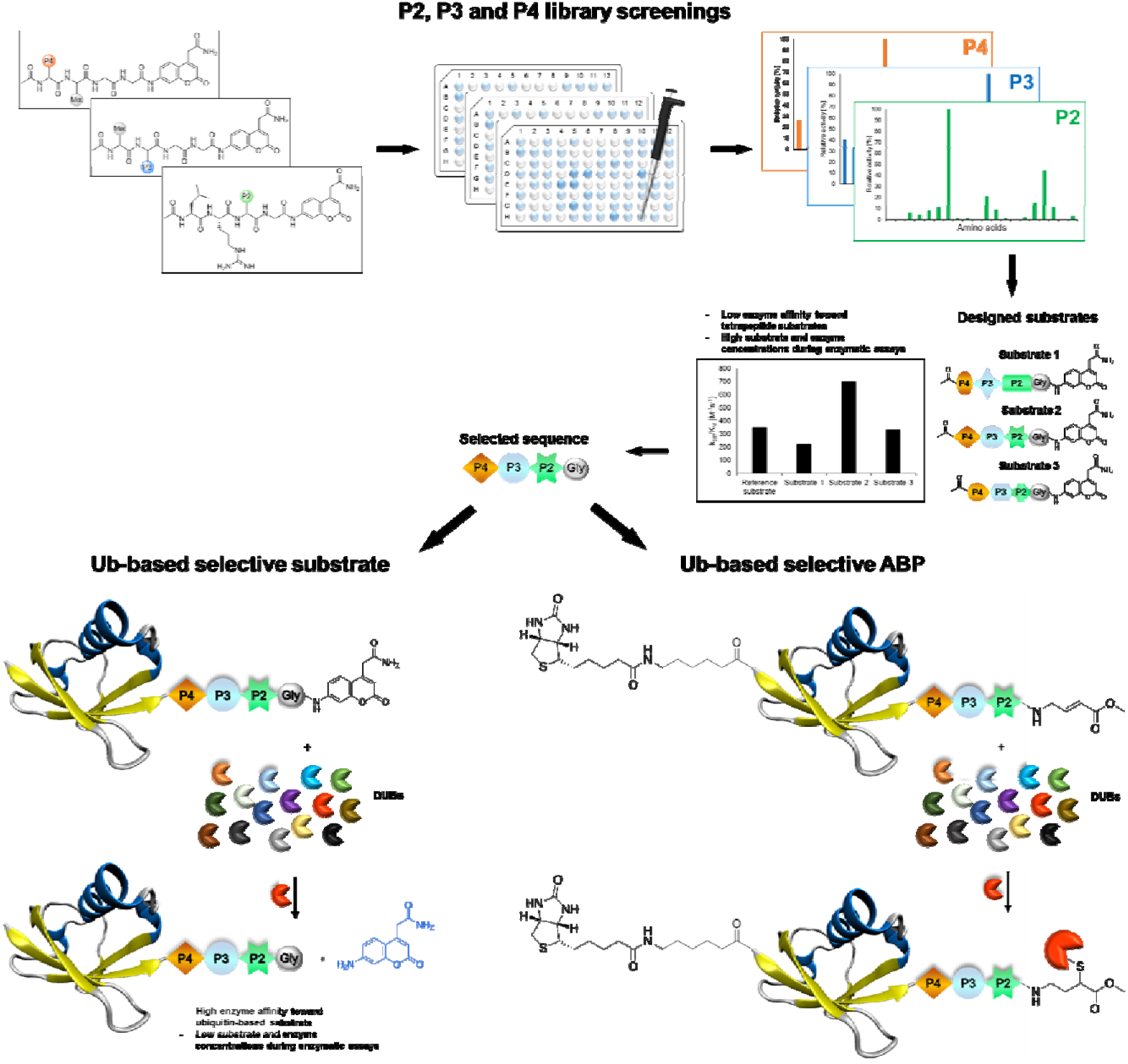
Workflow applied in the design of DUB-selective substrates and ABPs (PDB ID: 1UBQ).

## 2. Materials and Methods

### Reagents

All chemical reagents were obtained from commercial suppliers and used without further purification. The reagents used for the solid-phase peptide synthesis (SPPS) were as follows: Rink Amide (RA) resin (particle size 100-200 mesh, loading 0.74 mmol/g), 2-chlorotrityl chloride resin (particle size 100-200 mesh, loading 0.97 mmol/g), all Fmoc-amino acids, *O*-benzotriazole-*N,N,N‘,N‘*-tetramethyl-uronium-hexafluoro-phosphate (HBTU), 2-(1-H-7-azabenzotriazol-1-yl)-1,1,3,3-tetramethyluranium hexafluorophosphate (HATU), piperidine, diisopropylcarbodiimide (DICI) and trifluoroacetic acid (TFA), purchased from Iris Bioteh GmbH (Marktredwitz, Germany); Fmoc-Asp(O*t*Bu)-(Dmb)Gly-OH and Fmoc-Ser(*t*Bu)-Thr(ψ^Me,Me^pro)-OH (dipeptide building blocks) from AAPPTec (Louisville, KY, USA); 9-fluorenylmethyl carbazate (Fmoc-NH-NH_2_), Tris(2-carboxyethyl)phosphine hydrochloride (TCEP) and 2,2’-azobis[2-(imidazolin-2-yl)propane] (VA-044) from Combi-Blocks (San Diego, CA, USA); 4-mercaptophenylacetic acid (MPAA), guanidine hydrochloride (Gn*HCl), sodium phosphate monobasic monohydrate (NaH_2_PO_4_*H_2_O) and 4.0 M HCl in dioxane; 2-methyl-2-propanethiol (*t*BuSH) from Sigma-Aldrich (Poznan, Poland); NaNO_2_ from Merck Millipore (Warsaw, Poland); anhydrous *N*-hydroxybenzotriazole (HOBt) from Creosauls Louisville, KY, USA; 2,4,6-collidine (2,4,6-trimethylpyridine), HPLC-grade acetonitrile, and triisopropylsilane (TIPS) from Sigma-Aldrich (Poznan, Poland); and *N,N*-diisopropylethylamie (DIPEA) from VWR International (Gdansk, Poland). DMF *N,N*‘-dimethylformamide (DMF), dichloromethane (DCM), methanol (MeOH), diethyl ether (Et_2_O), acetic acid (AcOH), and phosphorus pentoxide (P_2_O_5_), obtained from Avantor (Gliwice, Poland). Individual substrates, Ub-based substrates and probes were purified by HPLC on a Waters M600 solvent delivery module with a Waters M2489 detector system using a semipreparative Wide Pore C8 Discovery column and Jupiter 10 µm C4 300 Å column (250 × 10 mm). The solvent composition was as follows: phase A (water/0.1% TFA) and phase B (acetonitrile/0.1% TFA). The purity of each compound was confirmed with an analytical HPLC system using a Jupiter 10 µm C4 300 Å column (250 × 4.6 mm). The solvent composition was as follows: phase A (water/0.1% TFA) and phase B (acetonitrile/0.1% TFA); gradient, from 5% B to 95% B over a period of 15 or 20 min. The molecular weight of each substrate and ABP was confirmed by high-resolution mass spectrometry on a High-Resolution Mass Spectrometer Waters LCT premier XE with electrospray ionization (ESI) and a time-of-flight (TOF) module.

### Combinatorial substrate library synthesis

A combinatorial tetrapeptide fluorogenic substrate library was synthesized on a solid support according to published protocols [29, 30]. The library consisted of two tetrapeptide sublibraries. Each of the sublibraries was synthesized separately, and the general synthetic procedure is described for the P3 sublibrary as an example. In the first step, Fmoc-ACC-OH (25 mmol, 2.5 eq) was attached to the RA resin (13.5 g) using coupling reagents: HOBt (25 mmol, 2.5 eq) and DICI (25 mmol, 2.5 eq) in DMF. After 24 h, the Fmoc protecting group was removed with 20% piperidine in DMF. In the next step, Fmoc-Gly-OH (25 mmol, 2.5 eq) was coupled using HATU (25 mmol, 2.5 eq) and 2,4,6-collidine (25 mmol, 2.5 eq) in DMF. Then, Fmoc-Gly-OH (25 mmol, 2.5 eq) was attached to the H_2_N-Gly-ACC-resin using HOBt and DICI as coupling reagents (25 mmol, 2.5 eq). After glycine coupling, the Fmoc group was removed (20% piperidine in DMF), and the resin was washed with DCM and MeOH and dried over P_2_O_5_. Then, the dried resin was divided into 138 portions. To each portion of the H_2_N-Gly-Gly-ACC-resin, natural or unnatural amino acids were attached, and the Fmoc protecting group was removed (20% piperidine in DMF). To provide equimolar substitution of each natural amino acid in the P4 position, an isokinetic mixture of Fmoc-protected amino acids was utilized. The last two steps of P3 sublibrary synthesis included N-terminal acetylation (solution of AcOH, HBTU and DIPEA) and cleavage of peptides from the resin using a TFA:H_2_O:TIPS (95%:2.5%:2.5%, %v/v/v) mixture. Finally, the sublibrary was precipitated in Et_2_O, dissolved in a mixture of acetonitrile and water, lyophilized and dissolved in biochemical grade DMSO at a concentration of 20 mM. The obtained sublibrary was used for kinetic studies without further purification. The P4 sublibrary was synthesized in the same manner.

### Synthesis of the defined P2 library and individual substrates

The defined P2 library (Ac-Leu-Arg-P2-Gly-ACC, where P2 represents 128 natural and unnatural amino acids) and individual tetrapeptide fluorogenic substrates were synthesized on a solid support using the SPPS method as previously described [29, 31]. Each substrate was purified by HPLC and analyzed using analytical HPLC and HRMS. The purity of each compound was ≥95%. The P2 library and individual substrates were dissolved at 10 mM in DMSO and stored at -80°C until use.

### Library and substrate screening

All screenings were performed using a spectrofluorometer (Molecular Devices Spectramax Gemini XPS) in 96-well plates (Corning). The release of ACC was monitored continuously for 40 min at the appropriate wavelength (λ_ex_ = 355 nm, λ_em_ = 460 nm). For the assay, 1 µL of substrate in DMSO was used with 99 µL of enzyme. The enzyme was incubated in assay buffer (MERS PLpro: 20 mM Tris, 150 mM NaCl, 5 mM DTT, pH 8.0 [18]; UCH-L3: 50 mM HEPES, 0.5 mM EDTA, 5 mM DTT, pH 7.5 [22]) for 30 min at 37°C before addition to the substrates on a plate. The final substrate concentration in each well during the assays was 200 µM for combinatorial P3 and P4 sublibraries, 100 µM for the Ac-Leu-Arg-P2-Gly-ACC library and 10 µM for individual substrates. The enzyme concentration was 2-3 µM for MERS PLpro and 6-20 µM for UCH-L3. Kinetic assays were repeated at least three times, and the results are presented as the mean values with standard deviations. The linear part of each progress curve was used to determine the substrate hydrolysis rate. Substrate specificity profiles were established by setting the highest value of relative fluorescence unit per second (RFU/s) from each library position as 100% and adjusting other values accordingly.

### Synthesis of Ub derivatives

Currently, different methods can be applied to obtain Ub and poly-Ub derivatives [6, 32-34]. We modified existing strategies to develop a new, versatile approach that enables the synthesis of Ub-based substrates and ABPs containing unnatural amino acids. We employed this method to synthesize four different Ub-ACC substrates and four corresponding N-terminally biotinylated ABPs with a C-terminal VME electrophile. Each substrate and probe was synthesized by applying a peptide hydrazide ligation strategy. Peptide hydrazide segments were synthesized by SPPS, purified by HPLC and assembled together in a stepwise manner. Then, we performed radical desulfurization and conjugated H_2_N-Gly-ACC to obtain Ub-ACC substrates or H_2_N-Gly-VME to obtain biotin-6-Ahx-Ub-VME ABPs. Herein, we present a general protocol for Ub derivative synthesis based on the example of Ub-ACC. We started the synthesis with Fmoc-hydrazide resin preparation based on a modified published method [35]. 2-CTC resin (0.97 mmol/g) was swelled for 20 min in fresh DCM and washed with 3x DCM. Fmoc-NH-NH_2_ (4 eq) was dissolved in DMF:DCM (8/1, *v/v*), and DIPEA (10 eq) was added. The mixture was added to the resin. The reaction was carried out at room temperature. After 1 h, DIPEA (5 eq) was added. Fmoc-NH-NH_2_ coupling was carried out for 4 h. The resin was washed with 3x DMF. End-capping of residual active 2-CTC resin was performed for 30 min by adding a capping mixture (DMF/MeOH/DIPEA, 3.5/0.5/0.2, *v/v/v*). After capping, Fmoc-NH-NH-2-CTC resin was washed with 3x DMF. Ub derivatives were assembled from 3 peptide hydrazide segments: Ub[1-27]-NH-NH_2_, Ub[28-45]-(A^28^C)-NH-NH_2_ and Ub[46-75]-(A^46^C)-NH-NH_2_. During the synthesis of ABPs, the Biot-6-Ahx-Ub[1-27]-NH-NH_2_ segment was used instead of Ub[1-27]-NH-NH_2_. Naturally occurring Ala residues on the N-termini of Ub[28-45]-(A^28^C)-NH-NH_2_ and Ub[46-75]-(A^46^C)-NH-NH_2_ were exchanged to Cys residues to enable subsequent native chemical ligation (NCL) reactions. Segments were synthesized manually in a straightforward manner using Fmoc-protected amino acids. The synthesis of Ub[46-75]-(A^46^C)-NH-NH_2_ required the use of dipeptide building blocks at Asp^52^Gly^53^ (Fmoc-Asp(O*t*Bu)- (Dmb)Gly-OH) and Ser^65^Thr^66^ (Fmoc-Ser(*t*Bu)-Thr(Ψ^Me,Me^pro)-OH), as reported elsewhere [32]. Fmoc-based SPPS of peptide hydrazides was performed in glass peptide synthesis vessels. Fmoc-NH-NH-2-CTC resin was swelled for 20 min in DCM, deprotected with 20% piperidine in DMF (two cycles: 5 and 25 min) and washed with 7x DMF. The first amino acid (2.5 eq) was dissolved in DMF with coupling reagents (2.5 eq HATU and 2.5 eq 2,4,6-collidine). The amino acid was preactivated for 30 s and added to the resin. The mixture was gently agitated for 24 h. After coupling, the resin was washed with 3x DMF, and the Fmoc groups were removed as described previously. A solution of the next amino acid (2.5 eq) with HATU (2.5 eq) and 2,4,6-collidine (2.5 eq) in DMF was added to the resin. After complete coupling of the second amino acid, Fmoc deprotection was carried out, and the following amino acids or dipeptide building blocks were coupled with the same coupling-deprotection cycles. The progress of the coupling and Fmoc deprotection of primary amine groups was monitored with the ninhydrin test. If needed, double coupling was performed. Biotin in the Biot-6-Ahx-Ub[1-27]-NH-NH_2_ segment was coupled in DMF/DMSO (1:1, *v/v*). After the last coupling step, the resin was washed with 3x DMF, 3x DCM, and 3x MeOH and dried over P_2_O_5_ overnight. A cleavage cocktail (8 mL, TFA/H_2_O/TIPS, 95/2.5/2.5, %*v/v/v*) was added to the dry resin. After 2 h, the cleavage mixture was filtered from the resin and collected. Peptides were precipitated with cold Et_2_O and dried. Crude peptide hydrazides were purified by preparative HPLC and lyophilized. NCL reactions were performed based on a published protocol [36]. Ub[1-27]-NH-NH_2_ (7.13 µmol) was dissolved in ligation buffer (1.5 mL, 6 M Gn*HCl, 0.2 M NaH_2_PO_4_, pH 3.0-3.1) and cooled in an ice-salt bath to - 15°C for 20 min with stirring. Acyl azide generation was performed at -15°C for 20 min by the addition of NaNO_2_ (10 eq). Then, 1 mL of a solution of Ub[28-45]-(A^28^C)-NH-NH_2_ (1 eq) and MPAA (100 eq) in ligation buffer (pH adjusted to 6.5) was added to Ub[1-27]-N_3_. The ice-salt bath was removed, and when the reaction mixture reached room temperature, the pH was slowly adjusted to 6.8. The progress of the reaction was monitored by LC-MS. After completion of the reaction, reducing agent (TCEP, 200 µL, 0.1 M in ligation buffer, pH 6.5) was added. The ligation product (Ub[1-45]-(A^28^C)-NH-NH_2_) was purified by semipreparative HPLC and lyophilized. Ligation of Ub[1-45]-(A^28^C)-NH-NH_2_ with Ub[46-75]-(A^46^C)-NH-NH_2_ was performed in the manner described above. Next, the product of the second ligation (Ub[1-75]- (A^28^C,A^46^C)-NH-NH_2_) was desulfurized based on a previously published protocol [6]. Ub[1-75]-(A^28^C,A^46^C)-NH-NH_2_ (14.1 mg, 1.39 µmol) was dissolved in desulfurization buffer (1.8 mL, 6 M Gn*HCl, 100 mM Na_2_HPO_4_, 375 mM TCEP, 37.5 mM VA-044, 150 mM tBuSH, pH 6.9). Desulfurization was performed for 4 h at 37°C. The progress of the reaction was monitored by LC-MS. Ub[1-75]-NH-NH_2_ was isolated by semipreparative HPLC and lyophilized. The last step of synthesis was H_2_N-Gly-ACC conjugation. TFA*H_2_N-Gly-ACC was synthesized on RA resin (0.74 mmol/g) and used without further purification. Ub[1-75]-NH-NH_2_ (6.7 mg) was dissolved in DMF:DMSO (0.75 mL, 2:1, *v/v*). The mixture was cooled in an ice-salt bath to -20°C for 20 min with stirring. NaNO_2_ (10 eq) in DMF:DMSO (50 µL, 2:1, *v/v*) was added. The pH of the reaction mixture was adjusted to 3.0-4.0 with 4 M HCl in dioxane solution. Acyl azide generation was carried out for 30 min in an ice-salt bath, and the progress of the reaction was monitored by LC-MS. TFA*H_2_N-Gly-ACC (20 eq) in DMF:DMSO (100 µL, 2:1, *v/v*) and DIPEA (20 eq) were added to the acyl azide, and the pH was adjusted to 8.0-9.0 with DIPEA. The reaction was performed at -20°C for 1 h. The progress of the reaction was monitored by LC-MS. The product was purified by analytical HPLC (purity >95%), lyophilized, and characterized by HRMS. Synthesis of ABPs was performed in the same manner. The warhead synthesis protocol was adopted from a published method [37]. tBu-N-allyl carbamate (500 mg, 3.2 µmol) was dissolved in 10 mL of anhydrous toluene. Methyl acrylate (580 µL, 6.4 µmol), dichlorophenylborane (42 µL, 0.32 µmol) and 2^nd^ generation Grubbs catalyst (50 mg) were added. The reaction was carried out under reflux at 40°C with stirring overnight. After 12 h, the solvent was removed, and the mixture was purified by column chromatography on silica gel (Hex/EtOAc 5:1). The crude product was obtained in the form of a yellowish oil. *t*Bu group deprotection was performed by adding TFA/DCM/TIPS (4.2 mL, 3/1/0.2, *v/v/v*) cleavage mixture for 45 min with stirring. TFA*H_2_N-Gly-VME was then crystallized in cold Et_2_O and stored at -20°C.

### Determination of kinetic parameters for tetrapeptide and Ub-based substrates

Kinetic studies were carried out in 96-well plates. Wells contained 20 µL of ACC-labeled substrate at eight different concentrations (0.1–75 µM) and 80 µL of enzyme in the same assay buffer as that described above (10 nM–2.5 µM MERS PLpro; 1 nM–6 µM UCH-L3). Substrate hydrolysis was measured for 30 min using the following wavelengths: λ_ex_ = 355 nm, λ_em_ = 460 nm. Each experiment was repeated at least three times. Kinetic parameters were calculated using GraphPad Prism software with the Michaelis-Menten equation. Due to the precipitation of tetrapeptide substrates at high concentrations, only the specificity constant (k_cat_/K_M_) was determined. When [S_0_]<<K_M,_ the plot of v_i_ (the initial velocities) versus [S_0_] yields a straight line with slope representing V_max_/K_M_, k_cat_/K_M_ = slope/E (E – total enzyme concentration).

### Determination of DUB inhibition by Ub-based probes

To assess DUB inhibition by biotinylated Ub-based probes, recombinant enzymes (50 nM) in assay buffer were incubated with eight different probe concentrations (0–3 µM) for 30 min at 37°C. Then, Ub-ACC (30 µM) was added to estimate residual DUB activity. The release of ACC was monitored for 30 min at the following wavelengths: λ_ex_ = 355 nm and λ_em_ = 460 nm.

### DUB labeling in cell lysates

A-431 cells were cultured in DMEM supplemented with 10% fetal bovine serum, 2 mM L-glutamine, and antibiotics (100 U/mL penicillin, 100 µg/mL streptomycin) in a humidified 5% CO_2_ atmosphere at 37°C. Approximately 1 200 000 cells were harvested and washed three times with PBS. The cell pellet was lysed in buffer containing 20 mM Tris, 150 mM NaCl, and 5 mM DTT, pH 8.0, using a sonicator. The cell lysate was centrifuged for 10 min, and the supernatant was collected. Twenty microliters of lysate was incubated with 80 µL of Ub-based probes at different concentrations for 30 min at 37°C. Then, 50 µL of 3x SDS/DTT was added, and the samples were boiled for 5 min at 95°C and resolved on 4-12% Bis-Tris Plus 12-well gels at 30 µL sample/well. Electrophoresis was performed at 200 V for 29 min. Next, the proteins were transferred to a nitrocellulose membrane (0.2 µm, Bio-Rad) for 60 min at 10 V. The membrane was blocked with 2% BSA in Tris-buffered saline with 0.1% (v/v) Tween 20 (TBS-T) for 60 min at RT. Biotinylated Ub-based probes were detected with a fluorescent streptavidin Alexa Fluor 647 conjugate (1:10 000) in TBS-T with 1% BSA, and UCH-L3 was detected with a mouse anti-human monoclonal IgG_1_ antibody (1:1000) and fluorescent goat anti-mouse (1:10 000) using an Azure Biosystems Sapphire^™^ Biomolecular Imager and Azure Spot Analysis Software.

## 3. Results

### DUB substrate specificity profile

To date, the substrate specificity profile of selected DUBs has been determined using only natural amino acids [18, 22]. It has been found that these enzymes can also recognize other amino acids at the P4-P2 positions, not only the C-terminal LRGG motif. These results suggest that dissection of the binding pocket architecture of DUBs can lead to the development of new chemical tools for DUB investigation with high specificity. To precisely examine the binding pocket preferences of selected enzymes, we used substrate libraries containing natural and a large number of unnatural amino acids with diverse chemical structures. Since all DUBs recognize leucine and arginine at the P4 and P3 positions, respectively, we synthesized a defined library with a general structure of Ac-Leu-Arg-P2-Gly-ACC (where P2 is a natural or unnatural amino acid) to determine the substrate specificity profile of DUBs at the P2 position. To examine substrate preferences at the P4 and P3 positions, we designed and synthesized a hybrid combinatorial substrate library (HyCoSuL). This library consists of two sublibraries: Ac-P4-Mix-Gly-Gly-ACC and Ac-Mix-P3-Gly-Gly-ACC (where P4 and P3 represent 19 natural and 119 unnatural amino acids, and Mix represents an equimolar mixture of natural amino acids). We incorporated glycine at the P1 and P2 positions because this amino acid is preferred by all DUBs in the S1 and S2 pockets. These two libraries can be used to determine the substrate specificity profile at the P4-P2 positions of all enzymes that display deubiquitinating activity.

### P2 position

DUBs exhibited very narrow substrate specificity at the P2 position (**Figure 2A**, see full substrate specificity profile in the supplemental information, **Figure S1**). MERS PLpro recognized only glycine at this position. The S2 pocket of UCH-L3 could accommodate not only glycine as the best hit but also, to a lesser extent, some aliphatic amino acids, such as Ala (47%), Val (46.5%), Abu (33.5%), Nle (25%), 2-Aoc (24%), and Tle (15%), and large hydrophobic amino acids, such as Nle(OBzl) (38%), Glu(OBzl) (31%), and Bpa (25%).

**Figure 2.**
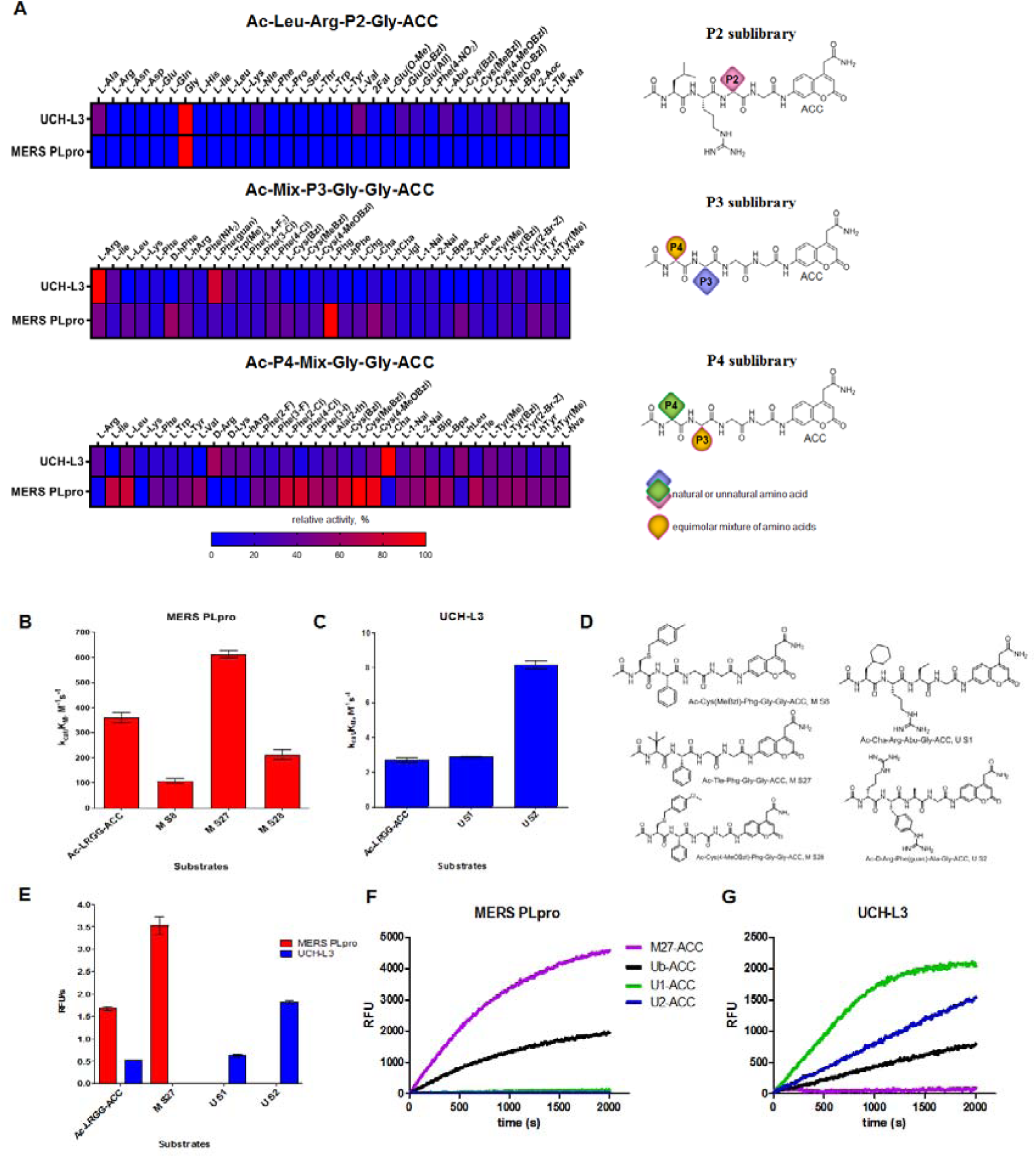
Substrate specificity of UCH-L3 and MERS PLpro. (A) Substrate specificity profiles of MERS PLpro and UCH-L3 presented as heat maps (left panel) and structures of libraries (right panel). Kinetic parameters for tetrapeptide fluorogenic substrates with MERS PLpro (B) and UCH-L3 (C). (D) Tetrapeptide substrate structures: M S8, Ac-Cys(MeBzl)-Phg-Gly-Gly-ACC; M S27, Ac-Tle-Phg-Gly-Gly-ACC; M S28, Ac-Cys(4-MeOBzl)-Phg-Gly-Gly-ACC; U S1, Ac-Cha-Arg-Abu-Gly-ACC; U S2, Ac-D*-*Arg-Phe(guan)-Ala-Gly-ACC. (E) Rate of tetrapeptide substrate hydrolysis by DUBs ([S]=10 µM; MERS PLpro concentration, 2.5 µM; UCH-L3 concentration, 6 µM); (F, G) Ub-based substrate selectivity ([S]=1 µM; MERS PLpro concentration, 5 nM; UCH-L3 concentration, 1 nM).

### P3 and P4 positions

UCH-L3 displayed a narrow substrate preference at the P3 position (**Figure 2A, S1**). Positively charged amino acids were preferred by the S3 pocket (Arg (100%), Phe(guan) (81%), and hArg (30%)). Hydrophobic residues were also recognized but with lower affinity (<30%, except Trp(Me) (41%) and Ile (36%)). In contrast, MERS PLpro preferred hydrophobic amino acids at the P3 position over basic residues (Phg (100%), D-hPhe (60%), Cha (54%), Arg (47%), 2-Aoc (44%), hArg (42%), hTyr (42%), and Leu (38.5%); amino acid structures presented in Table S2 supplemental information). The S4 pocket of UCH-L3 could accommodate hydrophobic amino acids (Cha (100%), Met (66.5%), hLeu (54%), 2-Nal (52.5%), Leu (46%), Cys(4-MeOBzl) (44%)) as well as positively charged L- and D-amino acids (D-Arg (64%), *D*-Lys (40.5%), hArg (39%), Arg (38%)). MERS PLpro favored aliphatic and bulky hydrophobic residues at the P4 position. The best recognized amino acids were Cys(MeBzl) (100%), Cys(4-MeOBzl) (93%), Cys(Bzl) (84%), Phe(4-Cl) (83%), Phe(2-Cl) (80%), Leu (79%), and Ile (75%). Basic and acidic residues were almost not recognized by MERS PLpro at this position.

### Kinetic analysis of tetrapeptide fluorogenic substrates

To select the optimal peptide sequence for MERS PLpro and UCH-L3, we analyzed substrate specificity preferences at the P4-P2 positions for each enzyme and selected amino acids that were both well recognized and selective toward the tested DUBs. For MERS PLpro, we chose Phg at the P3 position as the best recognized and most selective amino acid. At the P4 position, we selected Cys(MeBzl), Cys(4-MeOBzl) and Tle as the most promising candidates. The S2 pocket of UCH-L3 could accommodate some aliphatic amino acids; thus, we decided to incorporate Ala and Abu residues in substrate sequences. Although these amino acids were less well recognized than glycine, they can have a significant effect on substrate selectivity. At the P3 and P4 positions, we selected certain amino acids as the best hits (P3: Arg, Phe(guan); P4: D*-*Arg, Cha). After selection of the amino acids, we synthesized ACC-labeled substrates and determined the catalytic efficiency of the enzyme (**Figure 2B, C, D**). Most likely due to steric hindrance, Ac-Tle-Phg-Gly-Gly-ACC (M S27) was better hydrolyzed by MERS PLpro than the reference substrate (Ac-Leu-Arg-Gly-Gly-ACC). Ac-D*-*Arg-Phe(guan)-Ala-Gly-ACC (U S2) was 3 times better recognized by UCH-L3 than Ac-Leu-Arg-Gly-Gly-ACC. In the next step, we investigated substrate selectivity (**Figure 2E**). Kinetic analysis revealed that Ac-Tle-Phg-Gly-Gly-ACC was not recognized by UCH-L3, and Ac-D*-*Arg-Phe(guan)-Ala-Gly-ACC and Ac-Cha-Arg-Abu-Gly-ACC (U S1) were selective toward this protease. We selected these three peptide sequences for further analysis.

### Synthesis and kinetic analysis of fluorescent Ub derivative substrates

Since tetrapeptide fluorogenic substrates are not efficiently hydrolyzed by DUBs, even after incorporation of unnatural amino acids, the kinetic rates of the substrates were only 2-3 times higher compared to that of the reference Ac-LRGG-ACC substrate. Therefore, we decided to synthesize fluorescent Ub derivatives containing unnatural amino acids on the C-terminal tetrapeptidic epitope. To date, several chemical synthesis methods for Ub derivatives have been reported [6, 32-34]. We applied these methods with several modifications for effective synthesis of Ub-based substrates and ABPs comprising unnatural building blocks. In the first step, we divided Ub into three peptide segments and synthesized them separately using Fmoc-based SPPS of peptide hydrazides on 2-chlorotrityl chloride resin. This approach enables (1) efficient synthesis of each peptide segment with good isolated yields and purity; (2) incorporation of unnatural amino acids on the C-terminal Ub motif; and (3) modification of the N-terminus by introducing tags and linkers. In the next step, three peptide segments were assembled in a stepwise manner through NCL. Then, cysteine residues, which were introduced in the Ub sequence to enable NCL reactions, were converted to alanine using a free radical desulfurization reaction. In the last step, a fluorescent tag (ACC) with glycine was attached to the Ub[1-75]-NH-NH_2_ C-terminus, and the final product was purified and analyzed by HPLC and HRMS (**Figure 3**).

**Figure 3.**
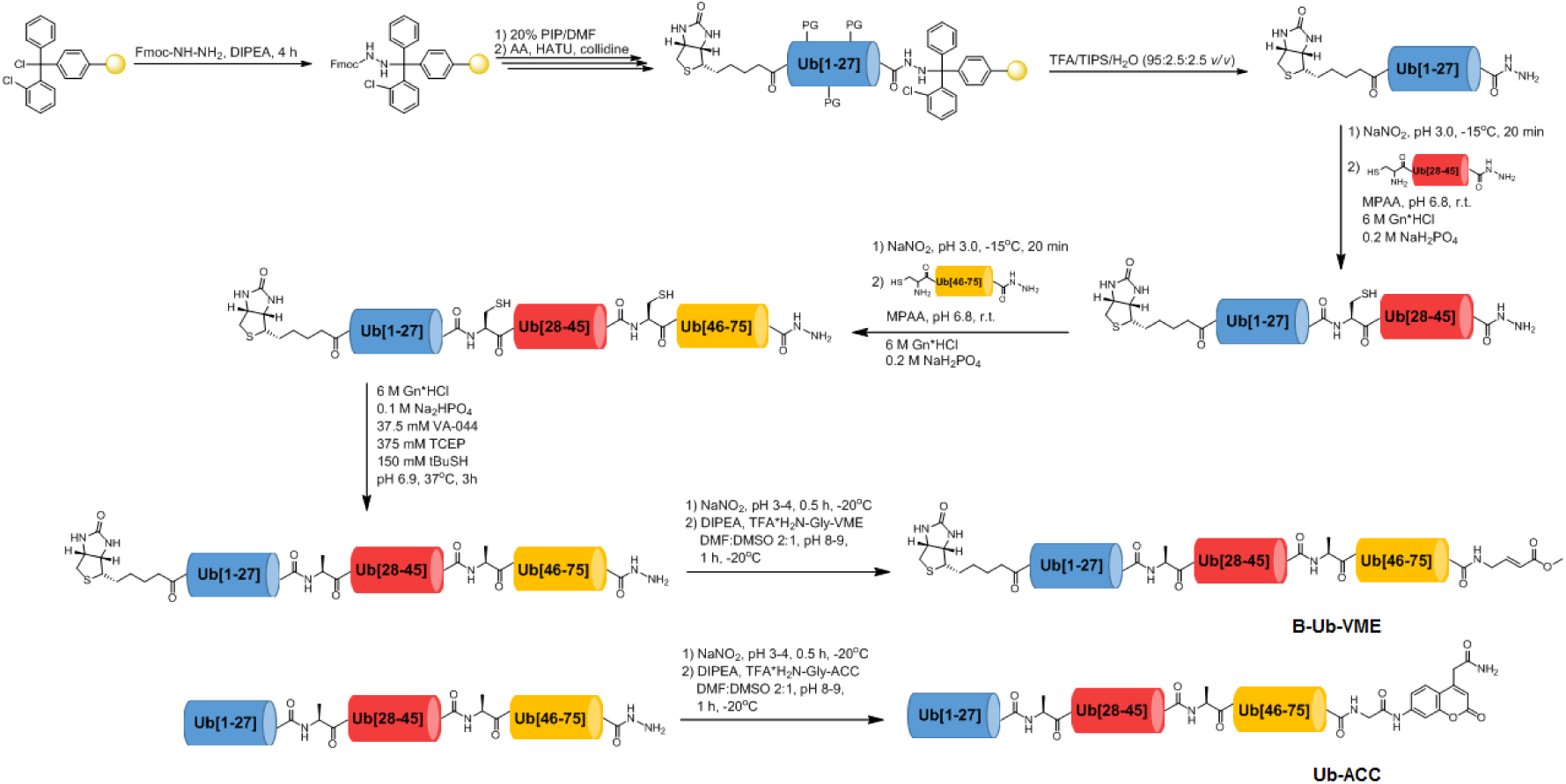
Scheme of Ub derivative synthesis.

Kinetic analysis of fluorescent Ub-based substrates revealed that by incorporation of unnatural amino acids in the Ub C-terminus, selective and active substrates for DUBs can be synthesized (**Figure 2F, G**). The designed substrates were better hydrolyzed by MERS PLpro (M27-ACC) and UCH-L3 (U1-ACC and U2-ACC) than Ub-ACC, used as a reference substrate (**Table 1**). U2-ACC was not recognized by MERS PLpro. M-ACC was poorly hydrolyzed by UCH-L3 (almost 200 times weaker than Ub-ACC).

**Table 1.** Kinetic parameters of Ub-based fluorogenic substrates with UCH-L3 and MERS PLpro.

| Ub-based substrate | Enzyme | $K_M$, $\mu M$ | $k_{cat}$, $s^{-1}$ | $k_{cat}/K_M$, $M^{-1}s^{-1}$ |
| --- | --- | --- | --- | --- |
| Ub-ACC | UCH-L3 | $0.048 \pm 0.002$ | $0.107 \pm 0.004$ | $(2.2 \pm 0.093) \times 10^6$ |
| | MERS PLpro | $3.55 \pm 0.09$ | $3.679 \pm 0.179$ | $(1.0 \pm 0.034) \times 10^6$ |
| M27-ACC | UCH-L3 | $2.11 \pm 0.07$ | $0.023 \pm 0.0005$ | $(1.1 \pm 0.060) \times 10^4$ |
| | MERS PLpro | $7.62 \pm 0.51$ | $12.386 \pm 0.341$ | $(1.6 \pm 0.076) \times 10^6$ |
| U1-ACC | UCH-L3 | $0.099 \pm 0.011$ | $0.387 \pm 0.017$ | $(3.9 \pm 0.22) \times 10^6$ |
|  | MERS PLpro | ND | ND | ND |
| U2-ACC | UCH-L3 | $0.083 \pm 0.005$ | $0.224 \pm 0.018$ | $(2.7 \pm 0.25) \times 10^6$ |
|  | MERS PLpro | - | - | - |

“ND” not determined due to substrate precipitation at high concentration
“-” not recognized

### Design of selective Ub-based probes

One of the main aims of this work was to design Ub-based probes for highly selective detection of the investigated DUBs. Thus, we converted our Ub-based substrates to probes by incorporating a detection tag on the N-terminus and an irreversible reactive group on the C-terminus. VME was applied as an electrophilic warhead due to its broad reactivity toward DUBs [11]. The biotin tag was separated from the peptide sequence by using a 6-aminohexanoic acid linker (6-Ahx). All Ub-based probes were synthesized using the same synthetic strategy as that used for the Ub-based substrates (**Figure 3**). The kinetic analysis with recombinant enzymes revealed that B-U2-VME was a potent and selective probe for UCH-L3 (**Table 2**). B-M27-VME was selective toward MERS PLpro (a much higher probe concentration was needed to inhibit UCH-L3). Furthermore, this probe was more potent toward MERS PLpro than the control probe B-Ub-VME (**Table 2**). The results were generally consistent with the Ub-based substrate kinetic data; however, B-U2-VME was a more potent probe for UCH-L3 than B-U1-VME.

**Table 2.** Kinetic analysis of DUB inhibition by Ub probes. Recombinant enzymes were incubated with various probe concentrations for 30 min at 37°C before DUB activity measurement. The experiment was performed in triplicate, and the average results are presented (SD<10%).

| Ub-probe,<br>[nM] | % enzyme inhibition ([E]=50 nM) |  |  |  |  |  |  |  |
| --- | --- | --- | --- | --- | --- | --- | --- | --- |
|  | UCH-L3 |  |  |  | MERS PLpro |  |  |  |
|  | B-Ub-VME | B-M27-VME | B-U1-VME | B-U2-VME | B-Ub-VME | B-M27-VME | B-U1-VME | B-U2-VME |
| 3000 | 97.8 | 65.2 | 49.8 | 87.5 | 100 | 100 | 71.5 | 0 |
| 1500 | 96.9 | 47.3 | 32.9 | 84.1 | 99.5 | 99.5 | 41.4 | 0 |
| 750 | 95.3 | 22.0 | 21.8 | 79.7 | 92.7 | 99.1 | 14.7 | 0 |
| 375 | 92.7 | 3.9 | 12.9 | 67.2 | 57.0 | 86.0 | 0 | 0 |
| 187.5 | 63.9 | 0 | 0 | 24.8 | 2.3 | 21.5 | 0 | 0 |
| 93.8 | 15.7 | 0 | 0 | 5.2 | 0 | 2.4 | 0 | 0 |
| 46.9 | 0 | 0 | 0 | 0 | 0 | 0 | 0 | 0 |
| 23.4 | 0 | 0 | 0 | 0 | 0 | 0 | 0 | 0 |

### Detection of DUBs in cell lysates

To assess Ub-based probe selectivity, we incubated cell lysates with various probe concentrations. Since UCH-L3 is expressed in the A-431 cell line, we selected this cell line for DUB labeling experiments. Due to the lack of MERS PLpro expression in human cell lines, we added the recombinant enzyme to cell lysates during incubation with probes. These DUB assays in cell lysates with designed probes revealed that our probes (B-M27-VME, B-U1-VME and B-U2-VME) displayed high selectivity toward MERS PLpro and UCH-L3 compared to the reference probe that labeled many cellular DUBs (**Figure 4A, B**). Labeling of UCH-L3 by our probes in A-431 cell lysate was confirmed by immunoblotting (**Figure 4C**).

**Figure 4.**
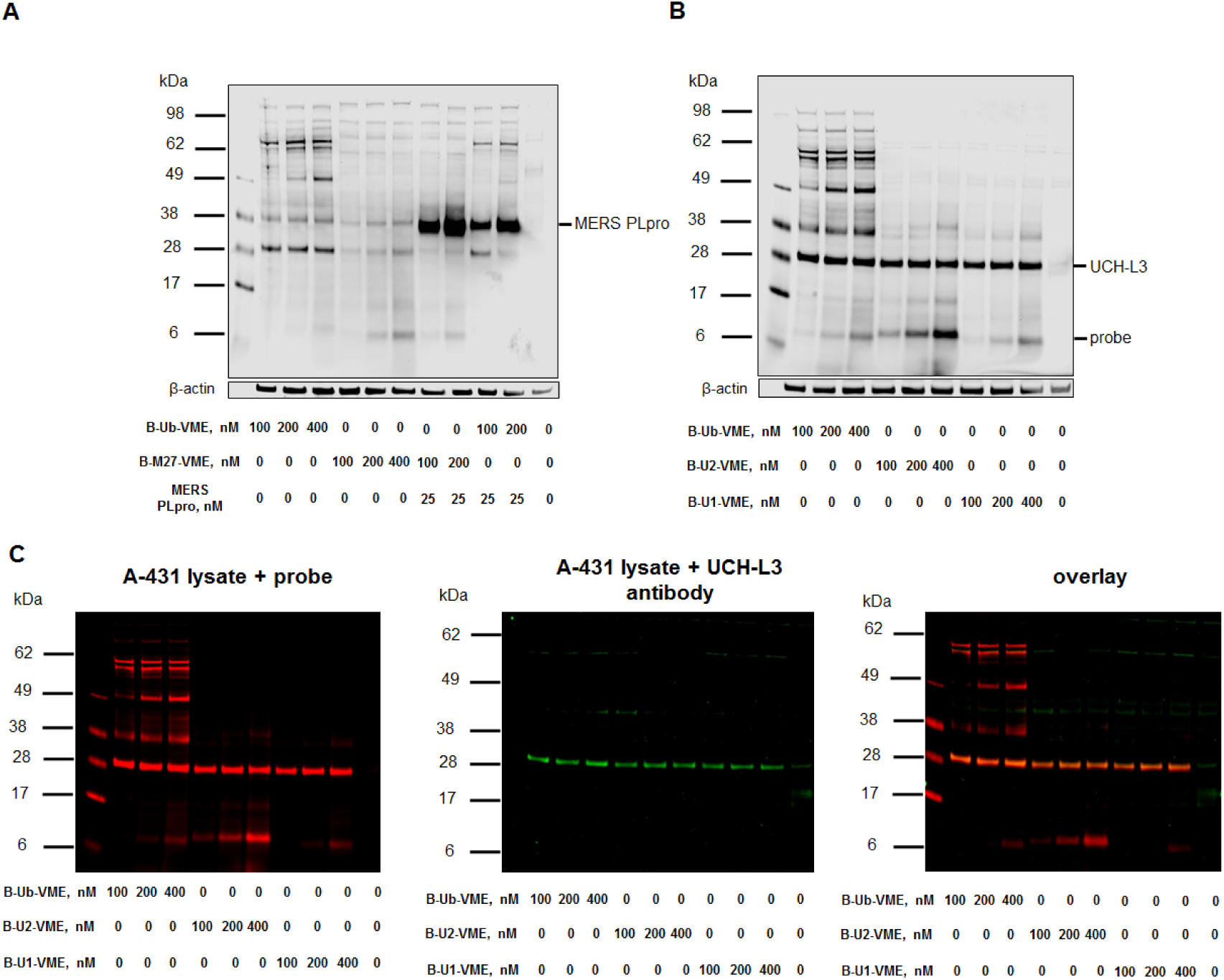
Ub-based probe selectivity. DUB labeling in cell lysates by B-Ub-VME and B-M27-VME (A) and B-U1-VME and B-U2-VME (B). (C) Detection of UCH-L3 in A-431 cell lysates using (1) Ub-based probes (using a streptavidin Alexa Fluor 647 conjugate) and (2) a UCH-L3 antibody. A-431 cell lysate was incubated with three different probe concentrations (100, 200 and 400 nM) for 30 min at 37°C. β-Actin was used as a loading control.

## 4. Discussion and Conclusions

DUBs constitute potential therapeutic targets in the treatment of viral diseases, neurodegenerative diseases and cancer [4, 14, 38]. The hottest example here is present influenza outbreak caused by Wuhan virus. Current studies indicate that this coronavirus is similar to SARS and MERS. Provided by us, this technology could be used for fast delivery of a selective 2019-nCoV PLpro DUB substrate and probe, which could effectively be used as a screen/reporter assay to find drugs that can specifically inhibit the 2019-nCoV PLpro DUB by screening infected patient cells without the need to purify DUB from infected patient cells. In classic examples in which DUBs are involved, enzymes belonging to the USP subclass play a key role in many biological processes and pathological conditions, including DNA repair, stabilization/degradation of the p53 protein, and progression of lung, breast, kidney and prostate cancer [39, 40]. UCH-L3 is overexpressed in triple-negative breast cancer (TNBC), which correlates with poor prognosis [27]. This enzyme is also involved in the progression of pancreatic cancer, but little is known about its precise functions [28]. This is due to the lack of selective chemical tools for the study of this enzyme; such tools would enable the determination of the role of this enzyme in normal and pathological states and could offer leading structures and chemical starting points for drug development. DUBs are highly specific toward Ub; thus, the chemical tools most commonly used to investigate DUBs are based on a Ub scaffold. These broad-spectrum tools are valuable in DUB profiling assays. However, substrates and ABPs recognized by one or a narrow subset of DUBs would enable more precise examination of this group of enzymes. To date, Ub-based probe selectivity toward one or a narrow subset of DUBs has been achieved by (1) selecting various C-terminal reactive electrophilic groups [11, 41]; (2) modifying the Ub scaffold length [42]; (3) synthesizing poly-Ub chains with different linkage topologies [15]; and (4) introducing various mutations in the Ub sequence [20, 21]. Our previous work showed that Ub-based substrate selectivity can be accomplished by modification of the C-terminal LRGG motif of Ub [22]. The introduction of natural amino acids other than canonical LRG at the P4-P2 positions in the Ub derivative provided selectivity but reduced DUB activity by 20-50 times compared to the natural analog.

Based on this finding, we decided to incorporate unnatural amino acids in the C-terminal motif of Ub to improve DUB activity and selectivity toward Ub-based substrates and ABPs. For precise analysis of the DUB binding pocket architecture, we utilized a combinatorial approach and synthesized three tetrapeptide fluorogenic substrate libraries. Due to the low tetrapeptide substrate hydrolysis rate of DUBs, three positions in the combinatorial library (P4 and P3 sublibraries) were fixed, and one contained an equimolar mixture of 19 amino acids. The P2 sublibrary was designed by fixing the P4, P3 and P1 positions as canonical amino acids present in Ub. The P2 position contained 128 natural and unnatural amino acids. This approach allowed us to achieve the highest possible substrate concentration in each position of the sublibrary during screening assays. We did not examine P1 substrate specificity because crystal structures of DUBs showed that this position can be occupied only by a glycine residue [13]. Library screening revealed that DUBs possess wide substrate specificity at the P4 and P3 positions. The DUBs recognized some unnatural amino acids even better than leucine and arginine at the P4-P3 positions. These results shed new light on DUB substrate preferences and the architecture of their binding sites. Kinetic analysis of tetrapeptide fluorogenic substrates revealed that the designed substrates with unnatural amino acids were selective and 2-3 times better recognized than the control substrate (Ac-LRGG-ACC); however, these substrates were still very poorly hydrolyzed. To improve DUB activity toward the designed substrates, we decided to synthesize unnatural Ub by incorporating the selected peptide sequences instead of the canonical LRGG sequence at the C-terminus and adding a fluorescent tag. DUBs possess two recognition regions that are required for effective Ub substrate hydrolysis [43]. The first region (being the secondary binding site or the exosite) interacts with the Ub surface, while the second region is the active center of the DUB, where the C-terminal LRGG motif is bound. Ub binding through the first region (distant from the active site cleft) leads to large conformational changes in DUB binding pockets that are required for effective catalysis of substrates [43]. Due to the nature of Ub binding by DUBs, we were expecting that translation from tetrapeptide substrates to full-length Ub sequences containing 2-3 unnatural amino acids on the C-terminus may result in a slight loss of selectivity. Despite the lack of hydrolysis of the M S27 tetrapeptide substrate by UCH-L3, kinetic analysis of the designed Ub-based substrates revealed that the MERS PLpro substrate (M27-ACC) was recognized by UCH-L3, but this recognition was almost 200 times weaker than that of the reference Ub-ACC. The UCH-L3 substrate (U2-ACC) was not hydrolyzed by MERS PLpro even at high concentrations. Furthermore, these substrates were more efficiently hydrolyzed by DUBs than Ub-ACC. Interestingly, U S1, an approximately 3-fold weaker UCH-L3 tetrapeptide substrate than U S2, after conversion to the U1-ACC Ub substrate, had an almost 1.5-fold higher kinetic efficiency constant than the U S2 analog U2-ACC. Nevertheless, both U1-ACC and U2-ACC displayed high selectivity toward UCH-L3 and exhibited kinetic parameters superior to those of Ub-ACC. Both of these features make these substrates very valuable alternatives to standard reagents used in studies of UCH-L3.

One of the most commonly used chemical tools in protease investigation is ABPs. Therefore, we converted our Ub-based fluorogenic substrates to Ub-based probes by replacing the ACC fluorophore with the VME electrophilic group and attaching a biotin tag to the N-terminus. Cell lysate assays confirmed the dramatically high selectivity of these engineered Ub-based probes toward the investigated enzymes, especially when compared to the reference probe B-Ub-VME, which labeled many cellular DUBs. These results indicate that Ub-based selective chemical tools for DUBs can be obtained by introducing unnatural amino acids into the C-terminal Ub motif.

In summary, we demonstrated that our approach can be successfully applied in the design and synthesis of selective mono-Ub substrates and ABPs. Precise analysis of DUB binding pocket architecture using defined and combinatorial libraries with natural and unnatural amino acids allowed us to extract key residues that were introduced into the C-terminal motif of Ub and provided selectivity toward the investigated enzymes. Our findings expand the knowledge of DUBs, as well as the existing ‘toolbox’ of Ub-based biochemical tools to study this group of enzymes. Moreover, the presented chemical approach may be beneficial for the development of a) new tools for the investigation of DUBs from distinct subclasses; b) more selective tools based on poly-Ub chains with different topologies; and c) new selective tools for studies of other Ub-like modifiers (UBLs), e.g., SUMO proteins, ISG15 or Nedd8, and their conjugation/deconjugation biochemical machinery.

## Acknowledgments

This project was supported by the National Science Center grant 2015/17/N/ST5/03072 (Preludium 9) in Poland (W.R.) and the “TEAM/2017-4/32” project, which is carried out within the TEAM program of the Foundation for Polish Science, cofinanced by the European Union under the European Regional Development Fund (M.D.). W.R. is a beneficiary of a START scholarship from the Foundation for Polish Science. NIH grants GM107257 (T.T.H.), ES025166 (T.T.H.) and F32GM100598 (M.B.) provided additional support for this work.

## Author contributions

M.D. conceptualized the study; M. D., W.R. and M.Ż. designed the research; W.R. and M.Ż. performed the research and collected data; S.J.S., M.B. and T.T.H. contributed enzymes; M.D., W.R. and M.Ż. analyzed and interpreted the data and wrote the manuscript; and S.J.S., M.B. and T.T.H. critically revised the manuscript.

