## Supplemental file for "Engineered unnatural ubiquitin for optimal detection of deubiquitinating enzymes"

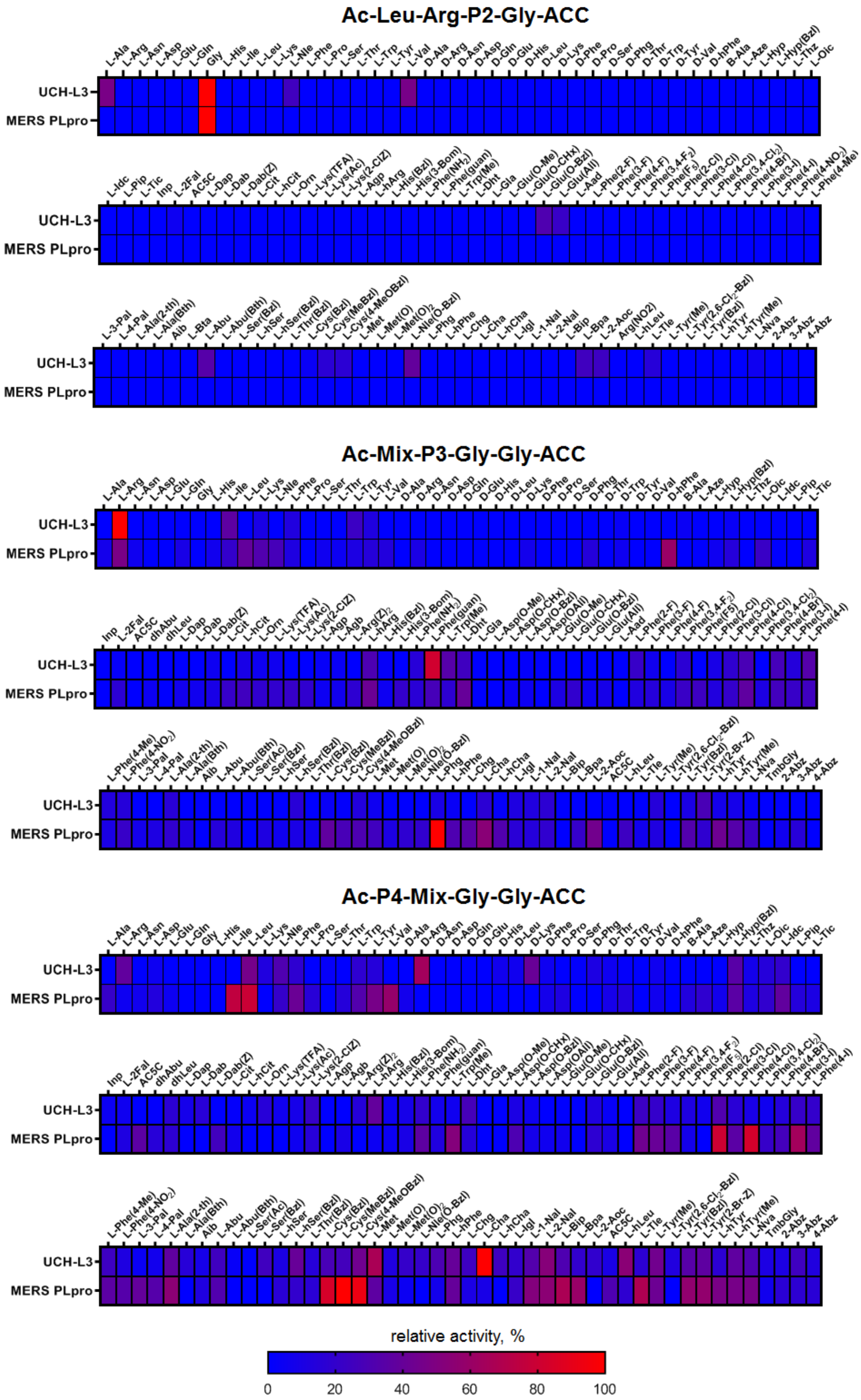


**Figure S1.** DUBs substrate specificity profiles presented as heat maps.

**Table S1.** Calculated and found m/z for Ac-Leu-Arg-P2-Gly-ACC substrate library.

| Substrate | Ac-Leu-Arg-P2-Gly-ACC | m/z calcd | m/z found |
|---|---|---|---|
| P2/1 | L-Ala | 658.3308 | 658.28 |
| P2/2 | L-Arg | 743.3948 | 743.37 |
| P2/3 | L-Asn | 701.3366 | 701.30 |
| P2/4 | L-Asp | 702.3206 | 702.28 |
| P2/5 | L-Glu | 716.3363 | 716.30 |
| P2/6 | L-Gln | 715.3522 | 715.15 |
| P2/7 | Gly | 644.3151 | 644.15 |
| P2/8 | L-His | 724.3526 | 724.25 |
| P2/9 | L-Ile | 700.3777 | 700.28 |
| P2/10 | L-Leu | 700.3777 | 700.29 |
| P2/11 | L-Lys | 715.3886 | 715.30 |
| P2/12 | L-Nle | 700.3777 | 700.20 |
| P2/13 | L-Phe | 734.3621 | 734.23 |
| P2/14 | L-Pro | 684.3464 | 684.19 |
| P2/15 | L-Ser | 674.3257 | 674.19 |
| P2/16 | L-Thr | 688.3413 | 688.21 |
| P2/17 | L-Trp | 773.3730 | 773.29 |
| P2/18 | L-Tyr | 750.3570 | 750.27 |
| P2/19 | L-Val | 686.3621 | 686.26 |
| P2/20 | D-Ala | 658.3308 | 658.20 |
| P2/21 | D-Arg | 743.3948 | 743.29 |
| P2/22 | D-Asn | 701.3366 | 701.24 |
| P2/23 | D-Asp | 702.3206 | 702.22 |
| P2/24 | D-Gln | 715.3522 | 715.25 |
| P2/25 | D-Glu | 716.3363 | 716.23 |
| P2/26 | D-His | 724.3526 | 724.27 |
| P2/27 | D-Leu | 700.3777 | 700.28 |
| P2/28 | D-Lys | 715.3886 | 715.29 |
| P2/29 | D-Phe | 734.3621 | 734.28 |
| P2/30 | D-Pro | 684.3464 | 684.22 |
| P2/31 | D-Ser | 674.3257 | 674.21 |
| P2/32 | D-Phg | 720.3464 | 720.25 |
| P2/33 | D-Thr | 688.3413 | 688.23 |
| P2/34 | D-Trp | 773.3730 | 773.31 |
| P2/35 | D-Tyr | 750.3570 | 750.30 |
| P2/36 | D-Val | 686.3621 | 686.25 |
| P2/37 | D-hPhe | 748.3777 | 748.29 |
| P2/38 | β-Ala | 658.3308 | 658.21 |
| P2/39 | L-Aze | 670.3308 | 670.23 |
| P2/40 | L-Hyp | 700.3413 | 700.22 |
| P2/41 | L-Hyp(Bzl) | 790.3883 | 790.23 |
| P2/42 | L-Thz | 702.3029 | 702.12 |
| P2/43 | L-Oic | 738.3934 | 738.23 |
| P2/44 | L-Idc | 732.3464 | 732.18 |
| P2/45 | L-Pip | 698.3621 | 698.19 |
| P2/46 | L-Tic | 746.3621 | 746.20 |

| | | | |
|---|---|---|---|
| P2/47 | Inp | 698.3621 | 698.20 |
| P2/48 | L-2Fal | 724.3413 | 724.17 |
| P2/49 | AC5C | 698.3621 | 698.19 |
| P2/50 | L-Dap | 673.3417 | 673.26 |
| P2/51 | L-Dab | 687.3573 | 687.26 |
| P2/52 | L-Dab(Z) | 821.3941 | 821.36 |
| P2/53 | L-Cit | 744.3788 | 744.32 |
| P2/54 | L-hCit | 758.3944 | 758.36 |
| P2/55 | L-Orn | 701.3730 | 701.29 |
| P2/56 | L-Lys(TFA) | 811.3709 | 811.32 |
| P2/57 | L-Lys(Ac) | 757.3992 | 757.28 |
| P2/58 | L-Lys(2-ClZ) | 883.3864 | 883.28 |
| P2/59 | L-Agp | 715.3635 | 715.24 |
| P2/60 | L-hArg | 757.4104 | 757.31 |
| P2/61 | L-His(Bzl) | 814.3995 | 814.31 |
| P2/62 | L-His(3-Bom) | 844.4101 | 844.34 |
| P2/63 | L-Phe($NH_2$) | 749.3730 | 749.29 |
| P2/64 | L-Phe(guan) | 791.3948 | 791.30 |
| P2/65 | L-Trp(Me) | 787.3886 | 787.29 |
| P2/66 | L-Dht | 775.3886 | 775.30 |
| P2/67 | L-Gla | 760.3261 | 760.23 |
| P2/68 | L-Glu(O-Me) | 730.3519 | 730.25 |
| P2/69 | L-Glu(O-cHx) | 798.4145 | 798.34 |
| P2/70 | L-Glu(O-Bzl) | 806.3832 | 806.32 |
| P2/71 | L-Glu(All) | 756.3676 | 756.31 |
| P2/72 | L-Aad | 730.3519 | 730.28 |
| P2/73 | L-Phe(2-F) | 752.3527 | 752.29 |
| P2/74 | L-Phe(3-F) | 752.3527 | 752.28 |
| P2/75 | L-Phe(4-F) | 752.3527 | 752.29 |
| P2/76 | L-Phe(3,4-$F_2$) | 770.3432 | 770.27 |
| P2/77 | L-Phe($F_5$) | 824.3150 | 824.26 |
| P2/78 | L-Phe(2-Cl) | 768.3231 | 768.24 |
| P2/79 | L-Phe(3-Cl) | 768.3231 | 768.25 |
| P2/80 | L-Phe(4-Cl) | 768.3231 | 768.25 |
| P2/81 | L-Phe(3,4-$Cl_2$) | 802.2841 | 802.31 |
| P2/82 | L-Phe(4-Br) | 812.2726 | 812.30 |
| P2/83 | L-Phe(3-I) | 860.2587 | 860.31 |
| P2/84 | L-Phe(4-I) | 860.2587 | 860.31 |
| P2/85 | L-Phe(4-Me) | 748.3777 | 748.40 |
| P2/86 | L-Phe(4-$NO_2$) | 779.3472 | 779.37 |
| P2/87 | L-3-Pal | 735.3573 | 735.35 |
| P2/88 | L-4-Pal | 735.3573 | 735.34 |
| P2/89 | L-Ala(2-th) | 740.3185 | 740.30 |
| P2/90 | L-Ala(Bth) | 791.3294 | 791.35 |
| P2/91 | Aib | 672.3464 | 672.30 |
| P2/92 | L-Bta | 790.3342 | 790.38 |
| P2/93 | L-Abu | 672.3464 | 672.32 |
| P2/94 | L-Abu(Bth) | 805.3451 | 805.41 |

| | | | |
|---|---|---|---|
| P2/95 | L-Ser(Bzl) | 764.3276 | 764.40 |
| P2/96 | L-hSer | 688.3413 | 688.31 |
| P2/97 | L-hSer(Bzl) | 778.3883 | 778.42 |
| P2/98 | L-Thr(Bzl) | 778.3883 | 778.43 |
| P2/99 | L-Cys(Bzl) | 780.3498 | 780.38 |
| P2/100 | L-Cys(MeBzl) | 794.3655 | 794.38 |
| P2/101 | L-Cys(4-MeOBzl) | 810.3604 | 810.33 |
| P2/102 | L-Met | 718.3342 | 718.27 |
| P2/103 | L-Met(O) | 734.3291 | 734.29 |
| P2/104 | L-Met$(O)_2$ | 750.3240 | 750.29 |
| P2/105 | L-Nle(O-Bzl) | 806.4196 | 806.40 |
| P2/106 | L-Phg | 720.3464 | 720.34 |
| P2/107 | L-hPhe | 748.3777 | 748.39 |
| P2/108 | L-Chg | 726.3934 | 726.39 |
| P2/109 | L-Cha | 740.4090 | 740.41 |
| P2/110 | L-hCha | 754.4247 | 754.44 |
| P2/111 | L-Igl | 760.3777 | 760.39 |
| P2/112 | L-1-Nal | 784.3777 | 784.42 |
| P2/113 | L-2-Nal | 784.3777 | 784.42 |
| P2/114 | L-Bip | 810.3934 | 810.44 |
| P2/115 | L-Bpa | 838.3883 | 838.44 |
| P2/116 | L-2-Aoc | 728.4090 | 728.40 |
| P2/117 | L-Arg$(NO_2)$ | 788.3799 | 788.40 |
| P2/118 | L-hLeu | 714.3934 | 714.37 |
| P2/119 | L-Tle | 700.3777 | 700.36 |
| P2/120 | L-Tyr(Me) | 764.3726 | 764.43 |
| P2/121 | L-Tyr(2,6-$Cl_2$-Bzl) | 908.3260 | 908.41 |
| P2/122 | L-Tyr(Bzl) | 840.4039 | 840.48 |
| P2/123 | L-hTyr | 764.3726 | 764.41 |
| P2/124 | L-hTyr(Me) | 778.3883 | 778.43 |
| P2/125 | L-Nva | 686.3621 | 686.33 |
| P2/126 | 2-Abz | 706.3308 | 706.33 |
| P2/127 | 3-Abz | 706.3308 | 706.31 |
| P2/128 | 4-Abz | 706.3308 | 706.30 |

**Table S2**. Structures of fixed natural and unnatural amino acids used in combinatiorial library.

| No | Structure + code | No | Structure + code | No | Structure + code |
|---|---|---|---|---|---|
| 1 | 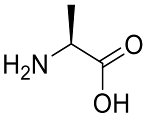 *L*-Ala | 2 | 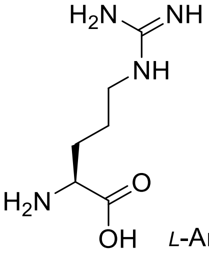 *L*-Arg | 3 | 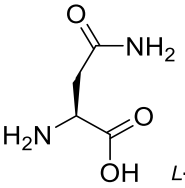 *L*-Asn |
| 4 | 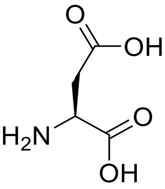 *L*-Asp | 5 | 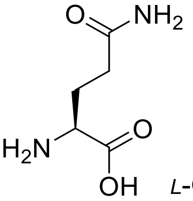 *L*-Gln | 6 | 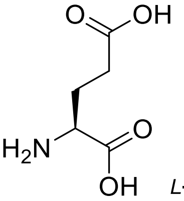 *L*-Glu |
| 7 | 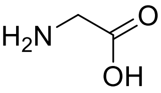 Gly | 8 | 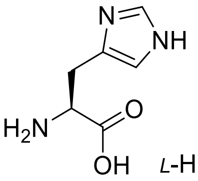 *L*-His | 9 | 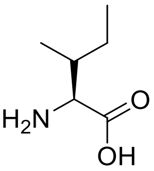 *L*-Ile |
| 10 | 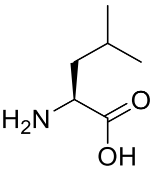 *L*-Leu | 11 | 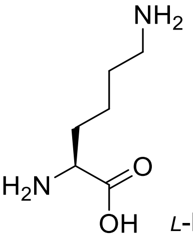 *L*-Lys | 12 | 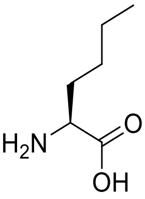 *L*-Nle |
| 13 | 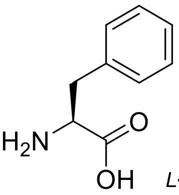 *L*-Phe | 14 | 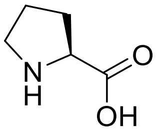 *L*-Pro | 15 | 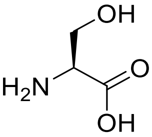 *L*-Ser |
| 16 | 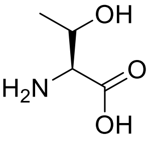 *L*-Thr | 17 | 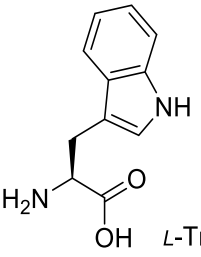 *L*-Trp | 18 | 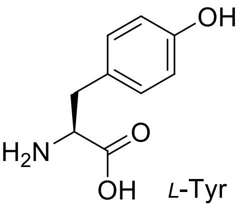 *L*-Tyr |

19

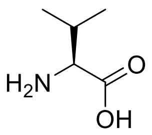

*L*-Val

20

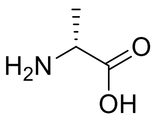

*D*-Ala

21

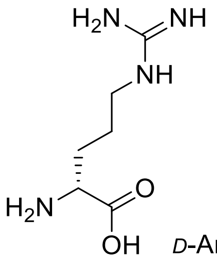

*D*-Arg

22

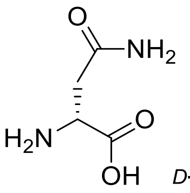

*D*-Asn

23

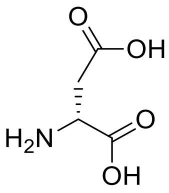

*D*-Asp

24

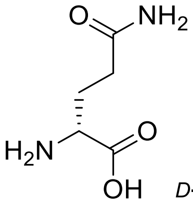

*D*-Gln

25

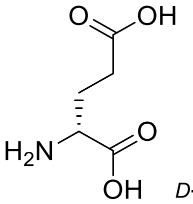

*D*-Glu

26

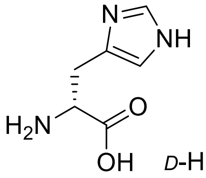

*D*-His

27

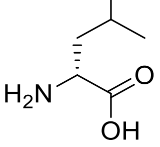

*D*-Leu

28

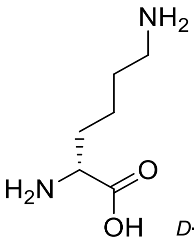

*D*-Lys

29

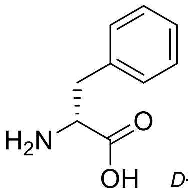

*D*-Phe

30

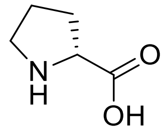

*D*-Pro

31

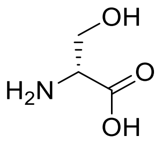

*D*-Ser

32

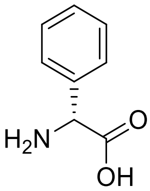

*D*-Phg

33

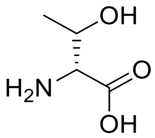

*D*-Thr

34

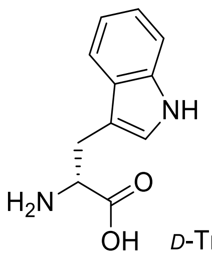

*D*-Trp

35

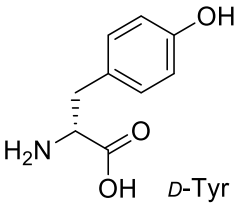

*D*-Tyr

36

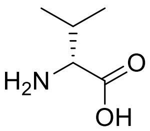

*D*-Val

37

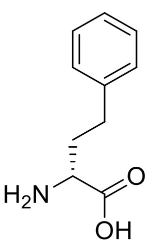

*D*-hPhe

38

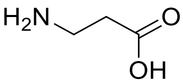

β–Ala

39

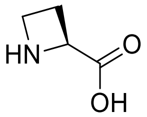

*L*-Aze

40 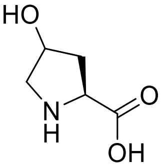 *L*-Hyp

41 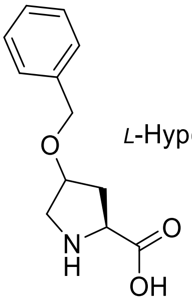 *L*-Hyp(Bzl)

42 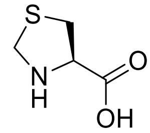 *L*-Thz

43 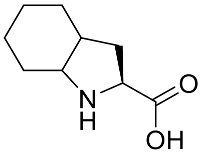 *L*-Oic

44 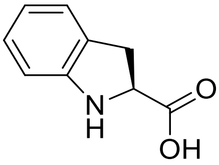 *L*-Idc

45 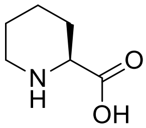 *L*-Pip

46 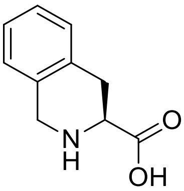 *L*-Tic

47 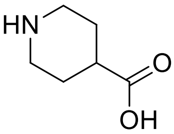 Inp

48 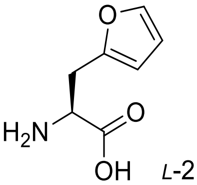 *L*-2Fal

49 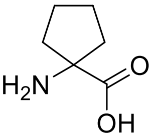 AC5C

50 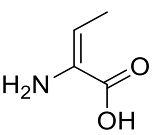 dhAbu

51 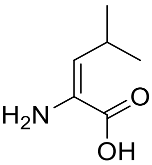 dhLeu

52 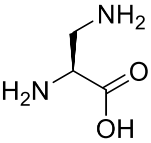 *L*-Dap

53 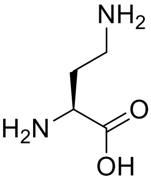 *L*-Dab

54 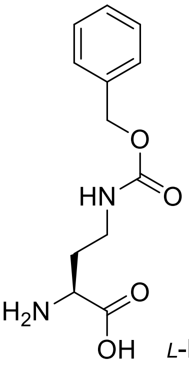 *L*-Dab(Z)

55 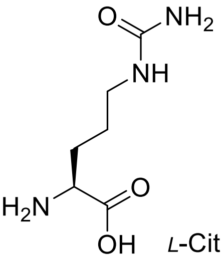 *L*-Cit

56 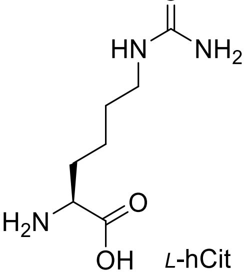 *L*-hCit

57 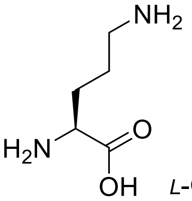 *L*-Orn

58 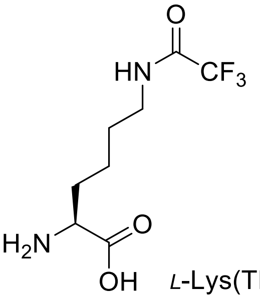 L-Lys(TFA)

59 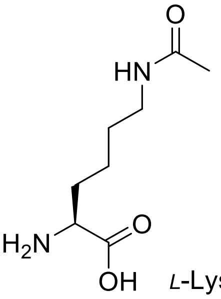 L-Lys(Ac)

60 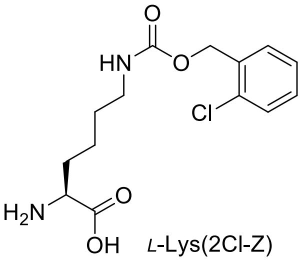 L-Lys(2Cl-Z)

61 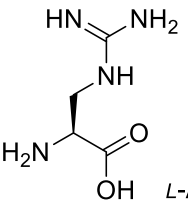 L-Agp

62 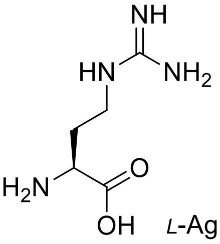 L-Agb

63 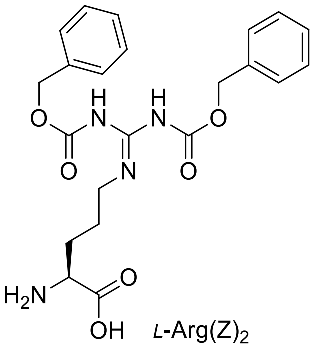 L-Arg(Z)$_2$

64 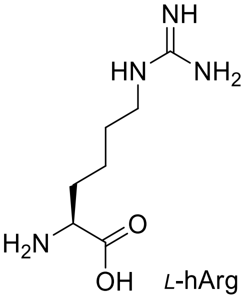 L-hArg

65 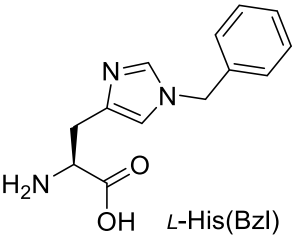 L-His(Bzl)

66 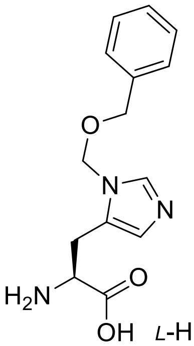 L-His(3-Bom)

67 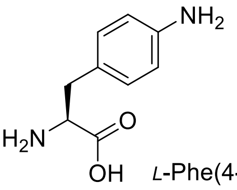 L-Phe(4-NH$_2$)

68 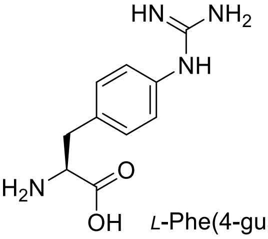 L-Phe(4-guan)

69 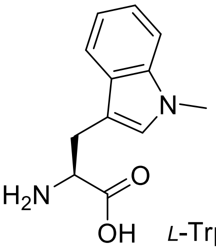 L-Trp(Me)

70 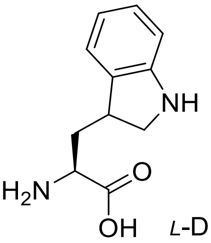 L-Dht

71 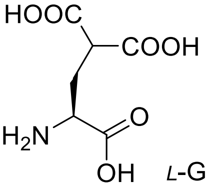 L-Gla

72 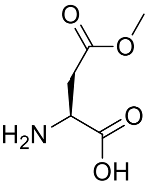 L-Asp(O-Me)

73 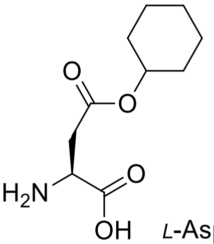 L-Asp(O-cHx)

74 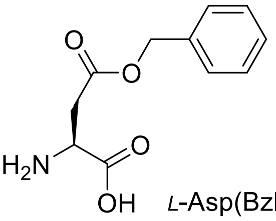 L-Asp(Bzl)

75 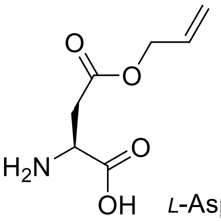 L-Asp(All)

76 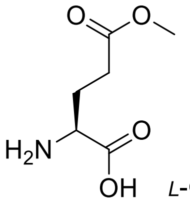 *L*-Glu(O-Me)

77 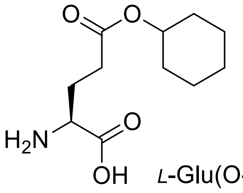 *L*-Glu(O-cHx)

78 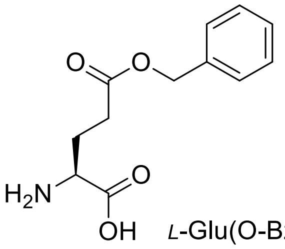 *L*-Glu(O-Bzl)

79 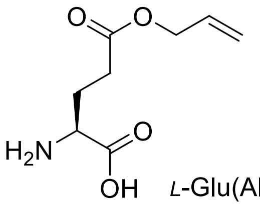 *L*-Glu(All)

80 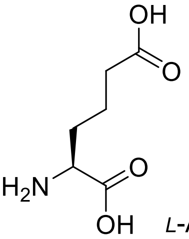 *L*-Aad

81 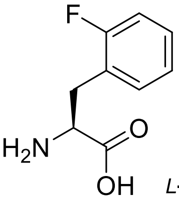 *L*-Phe(2-F)

82 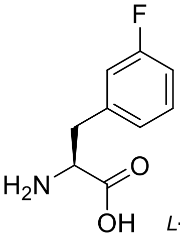 *L*-Phe(3-F)

83 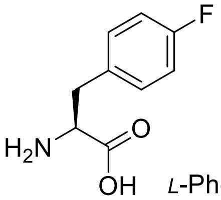 *L*-Phe(4-F)

84 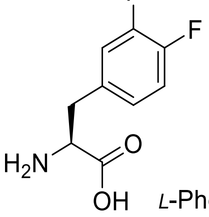 *L*-Phe(4,3-$F_2$)

85 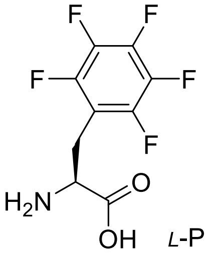 *L*-Phe($F_5$)

86 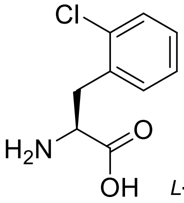 *L*-Phe(2-Cl)

87 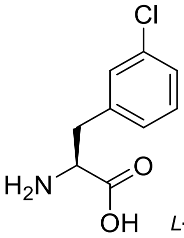 *L*-Phe(3-Cl)

88 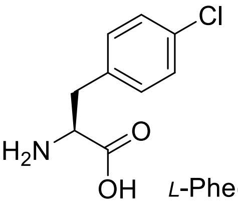 *L*-Phe(4-Cl)

89 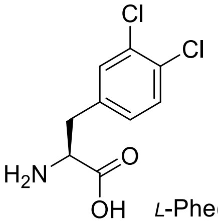 *L*-Phe(3,4-$Cl_2$)

90 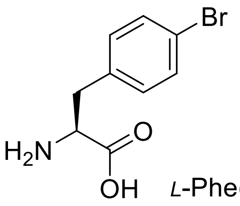 *L*-Phe(4-Br)

91 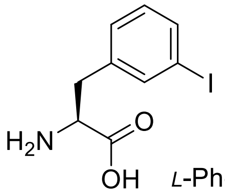 *L*-Phe(3-I)

92 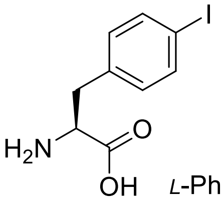 *L*-Phe(4-I)

93 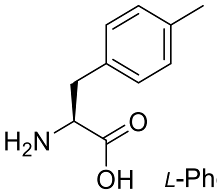 *L*-Phe(4-Me)

94 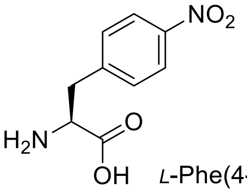 *L*-Phe(4-$NO_2$)

95 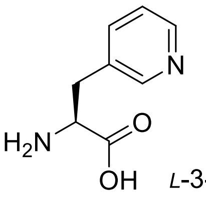 *L*-3-Pal

96 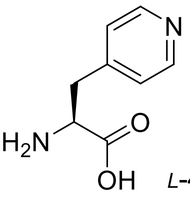 *L*-4-Pal

97 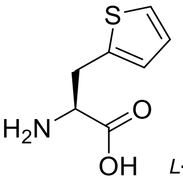 *L*-Ala(2-th)

98 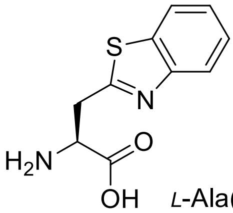 *L*-Ala(Bth)

99 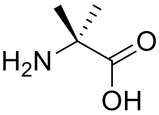 Aib

100 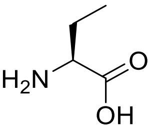 *L*-Abu

101 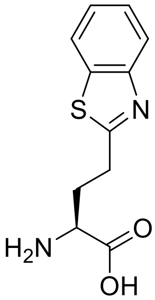 *L*-Abu(Bth)

102 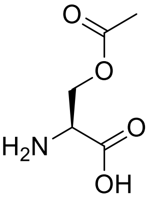 *L*-Ser(Ac)

103 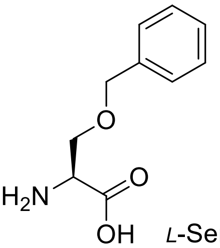 *L*-Ser(Bzl)

104 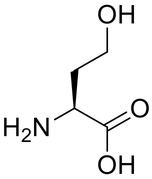 *L*-hSer

105 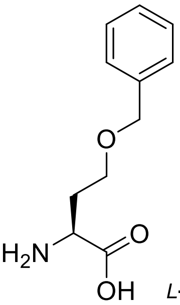 *L*-hSer(Bzl)

106 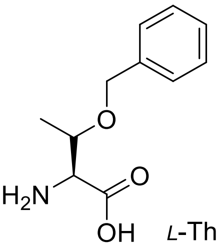 *L*-Thr(Bzl)

107 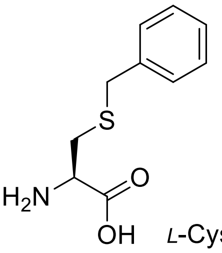 *L*-Cys(Bzl)

108 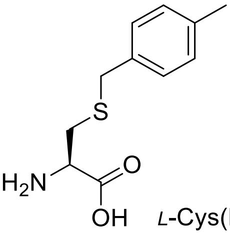 *L*-Cys(MeBzl)

109 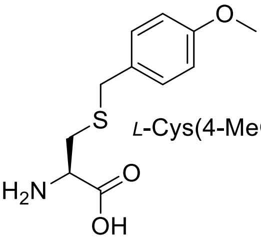 *L*-Cys(4-MeOBzl)

110 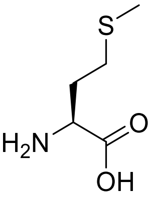 *L*-Met

111 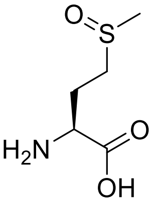 *L*-Met(O)

112 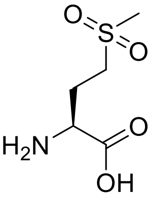 *L*-Met($O_2$)

113 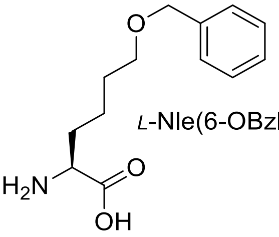 *L*-Nle(6-OBzl)

114 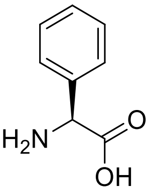 *L*-Phg

115

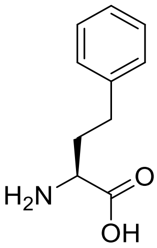

*L*-hPhe

116

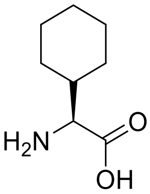

*L*-Chg

117

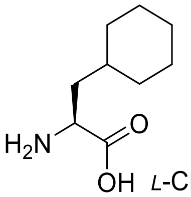

*L*-Cha

118

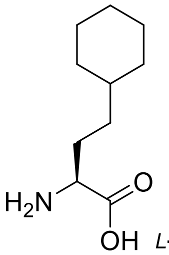

*L*-hCha

119

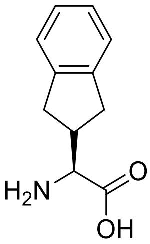

*L*-Igl

120

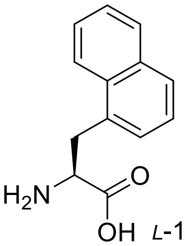

*L*-1-Nal

121

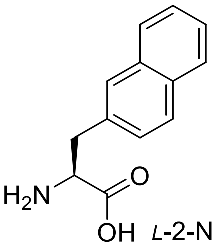

*L*-2-Nal

122

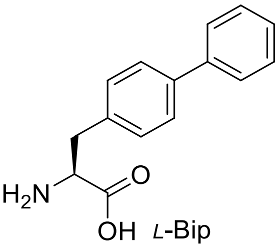

*L*-Bip

123

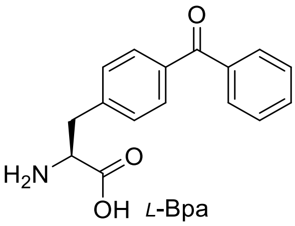

*L*-Bpa

124

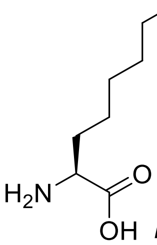

*L*-2-Aoc

125

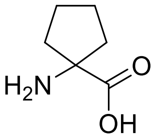

AC5C

126

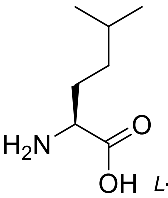

*L*-hLeu

127

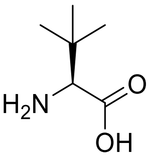

*L*-Tle

128

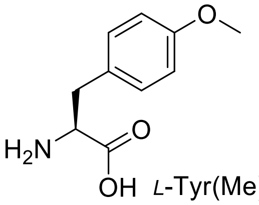

*L*-Tyr(Me)

129

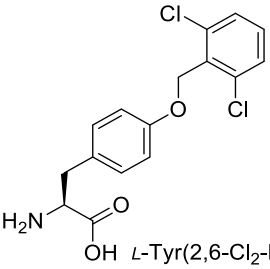

*L*-Tyr(2,6-$Cl_2$-Bzl)

130

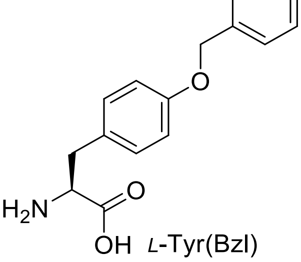

*L*-Tyr(Bzl)

131

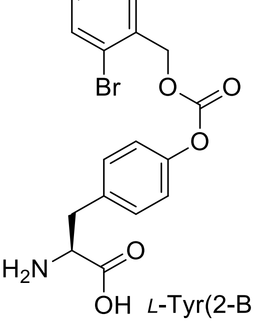

*L*-Tyr(2-Br-Z)

132

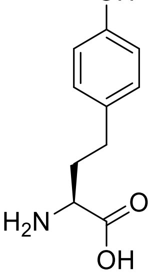

*L*-hTyr

133 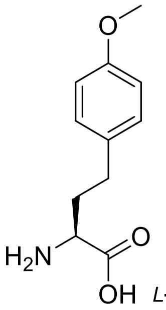 *L*-hTyr(Me)

134 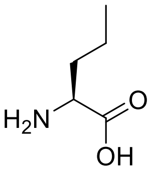 *L*-Nva

135 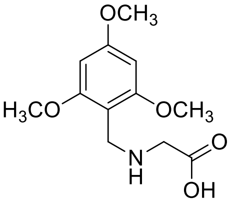 TmbGly

136 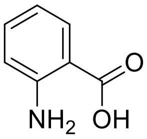 2-Abz

137 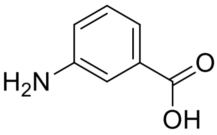 3-Abz

138 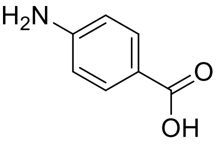 4-Abz

### HRMS and analytical chromatograms of synthesized substrates and activity based probes

#### Ac-Leu-Arg-Gly-Gly-ACC

HRMS for $C_{29}H_{41}N_9O_8$ (m/z $_{calcd}$ = 644.3151; m/z $_{found}$ = 644.3158)

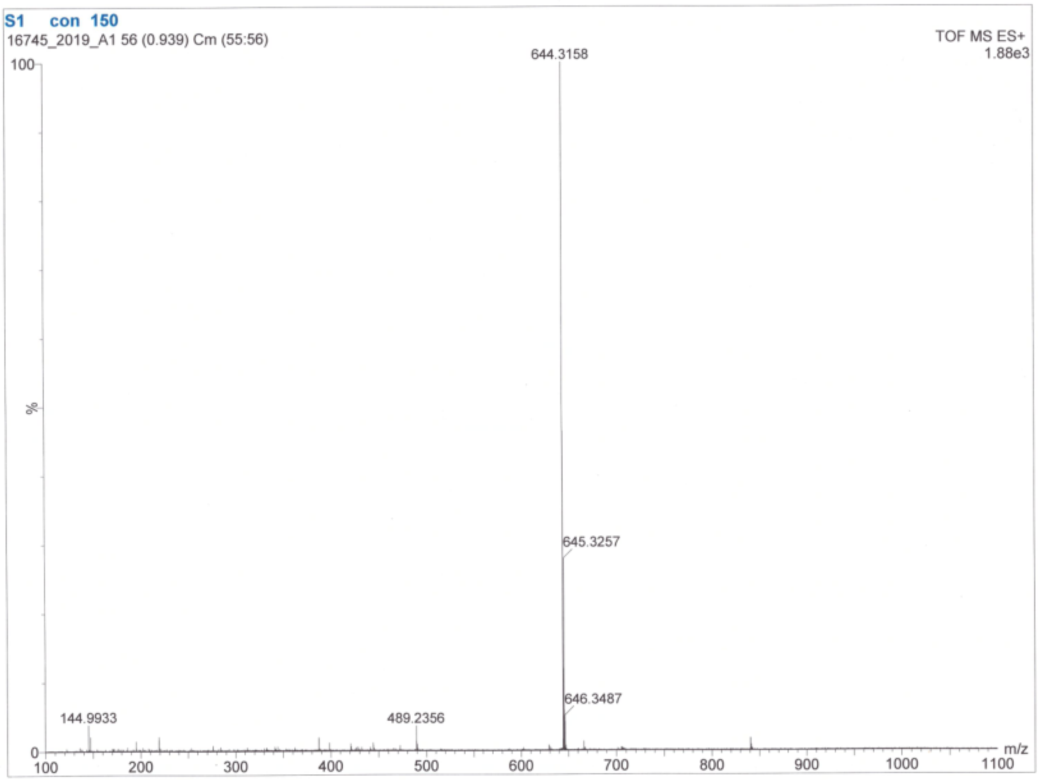


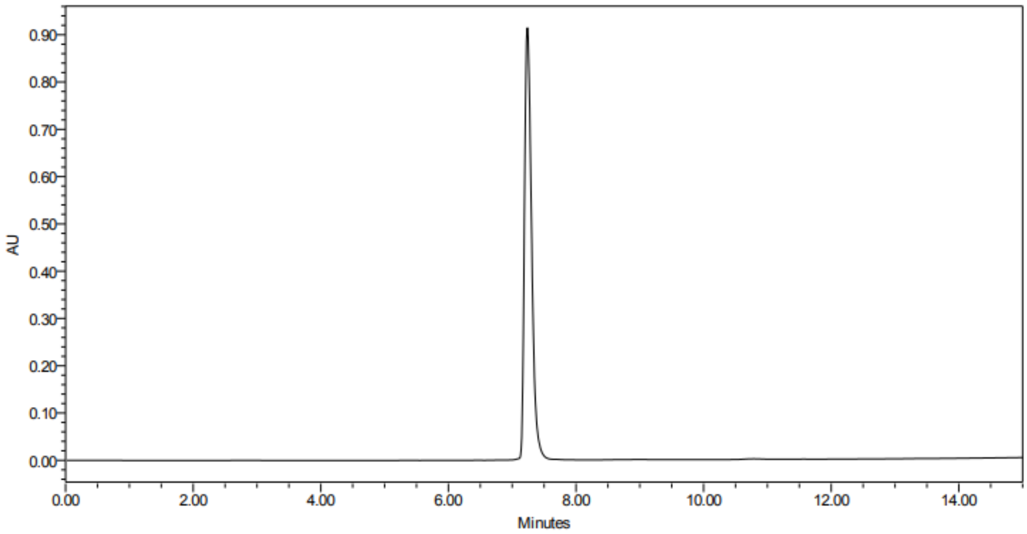

**Ac-Tle-Phg-Gly-Gly-ACC**

HRMS for $C_{31}H_{36}N_6O_8$ (m/z $_{calcd}$ = 621.2595; m/z $_{found}$ = 621.2672)

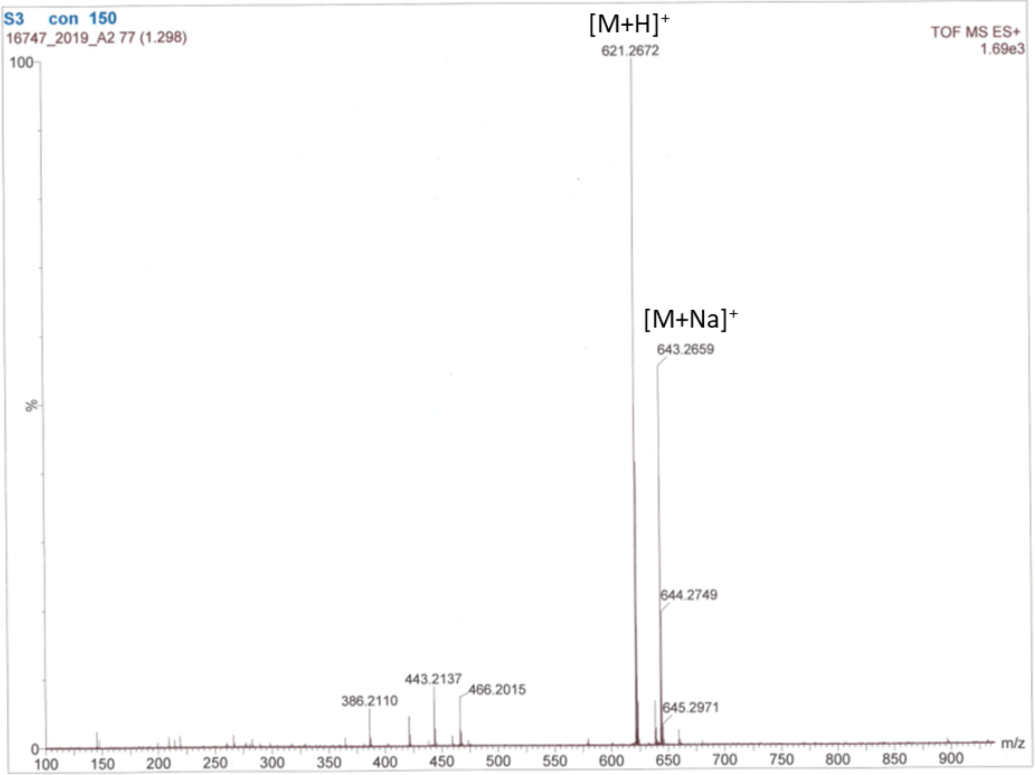


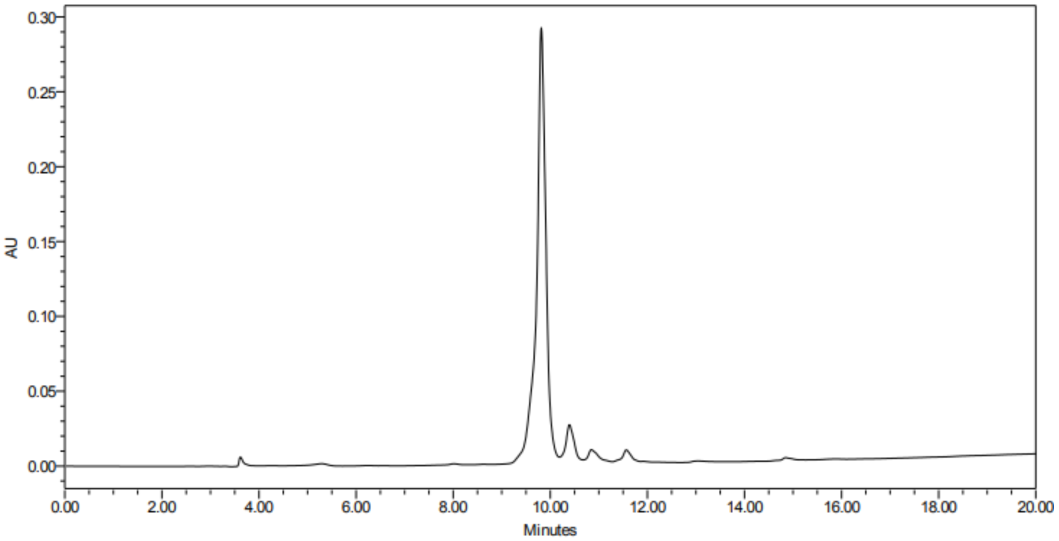

**Ac-Cha-Arg-Abu-Gly-ACC**

HRMS for $C_{34}H_{49}N_9O_8$ (m/z $_{calcd}$ = 712.3777; m/z $_{found}$ = 712.3780)

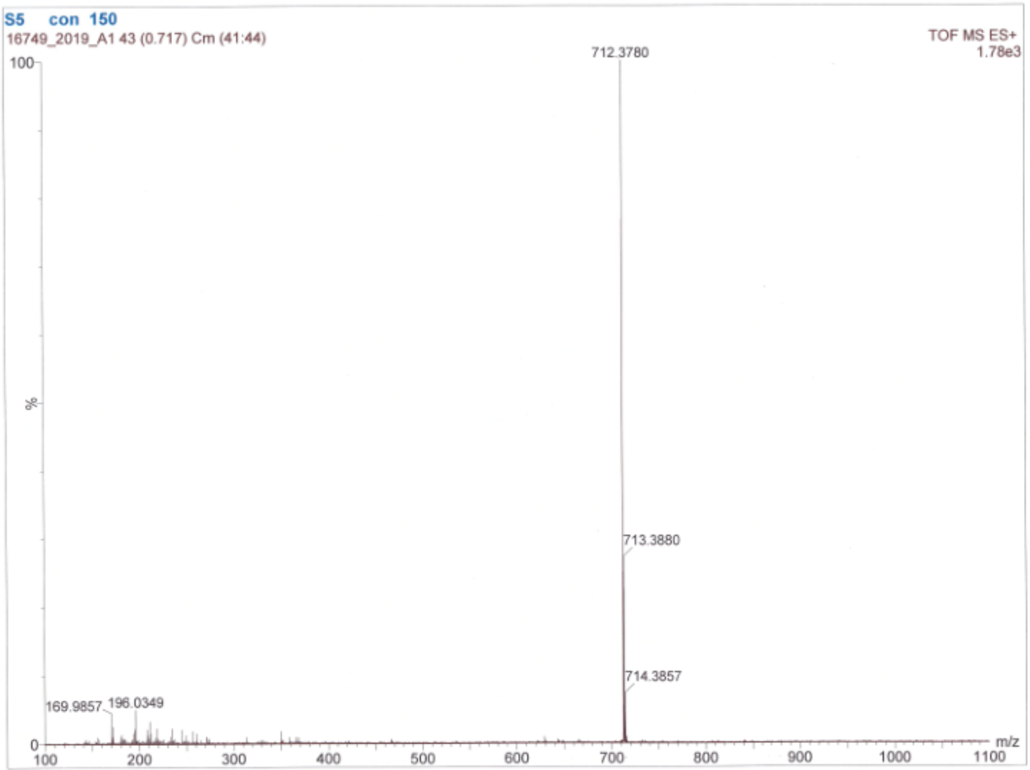


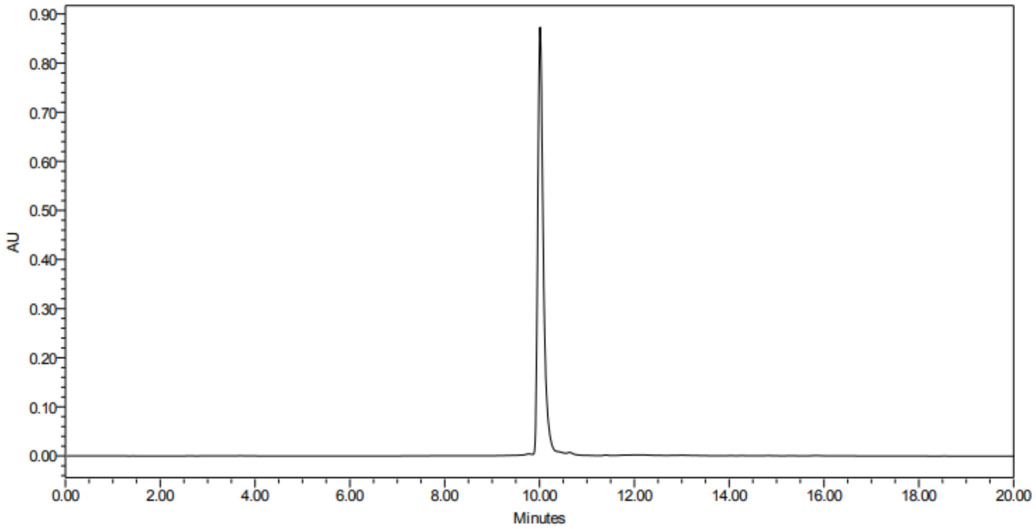

**Ac-DArg-Phe(guan)-Ala-Gly-ACC**

HRMS for $C_{34}H_{44}N_{12}O_8$ (m/z $_{calcd}$ = 749.3478; m/z $_{found}$ = 749.3479)

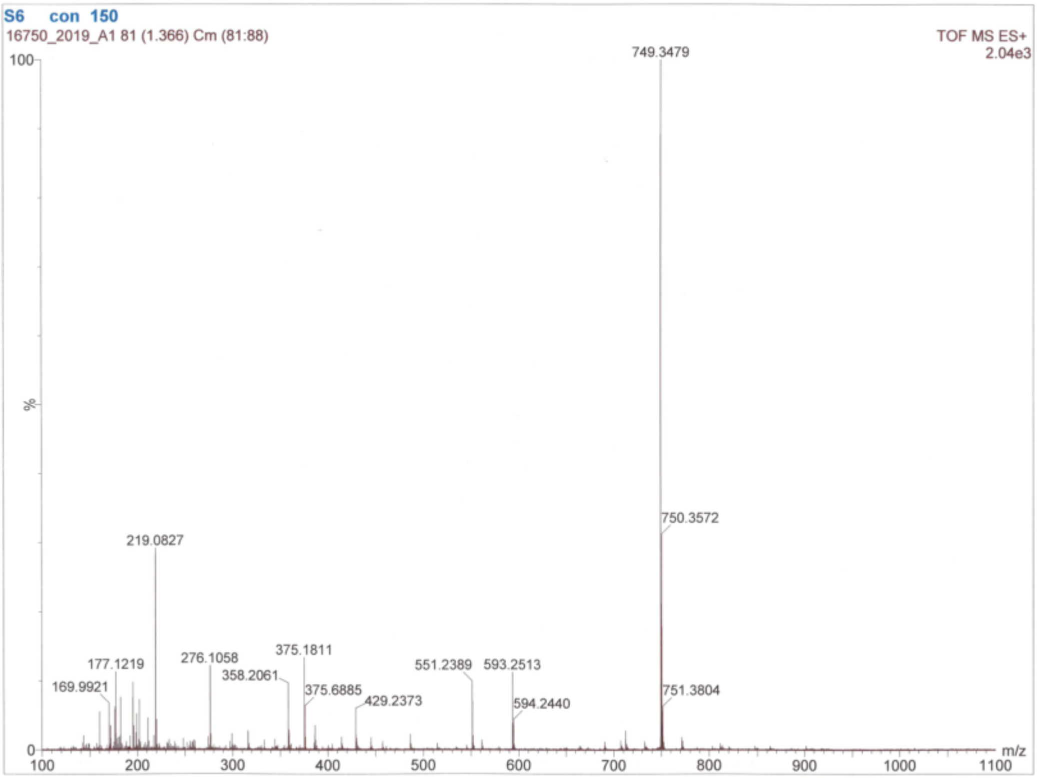


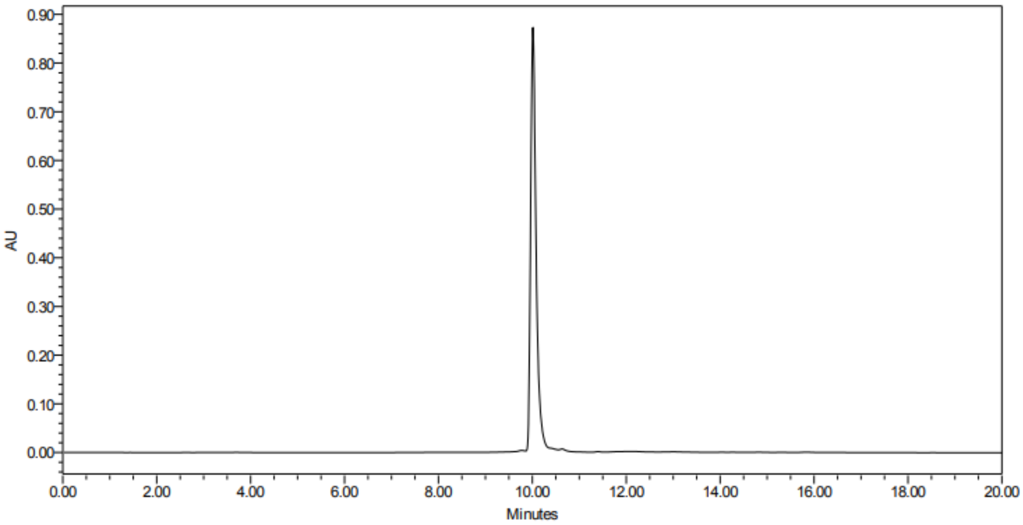

**Ac-Cys(4-MeOBzl)-Phg-Gly-Gly-ACC**

HRMS for $C_{36}H_{38}N_6O_9S$ (m/z $_{calcd}$ = 731.2494; m/z $_{found}$ = 731.2478)

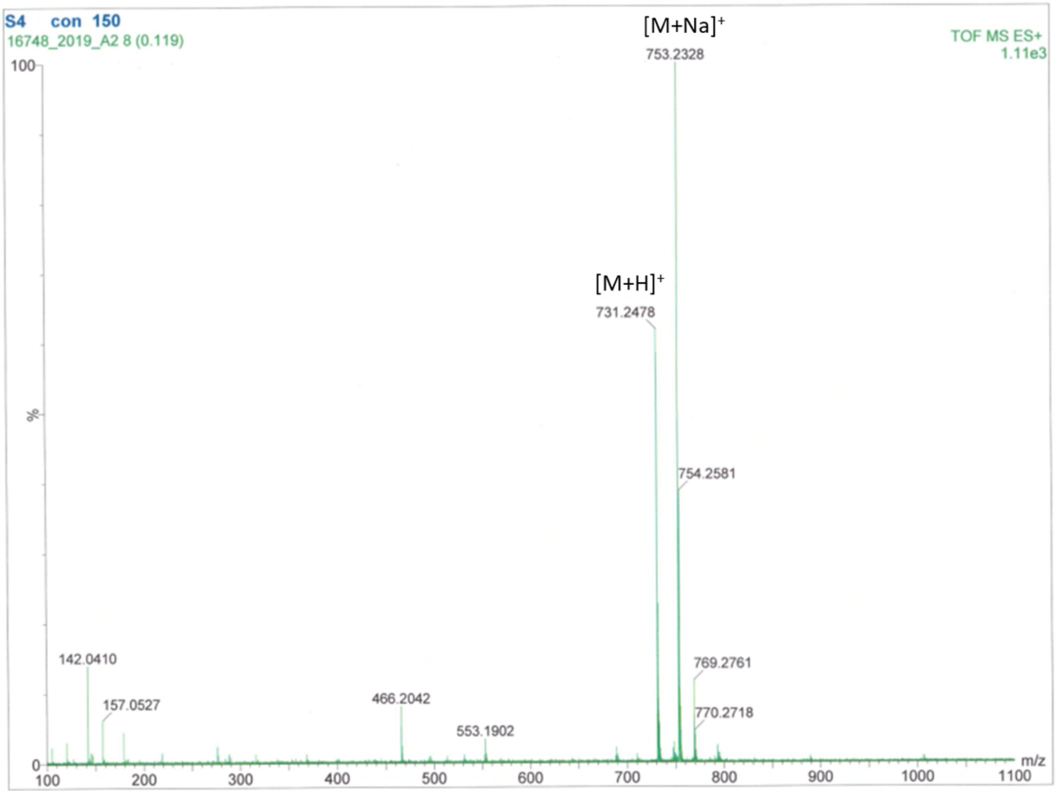


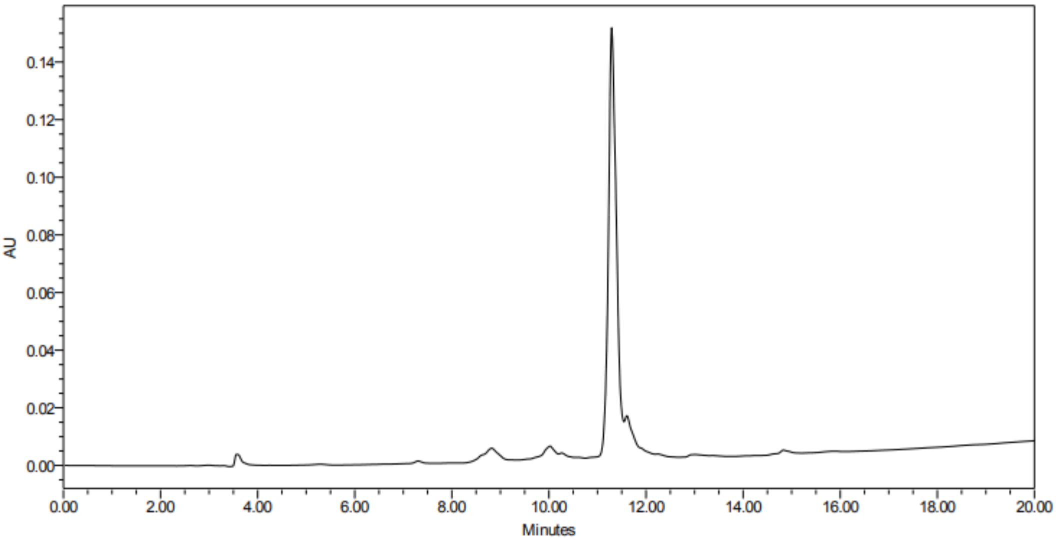

**Ac-Cys(MeBzl)-Phg-Gly-Gly-ACC**

HRMS for $C_{36}H_{38}N_6O_8S$ (m/z $_{calcd}$ = 715.2545; m/z $_{found}$ = 715.2556)

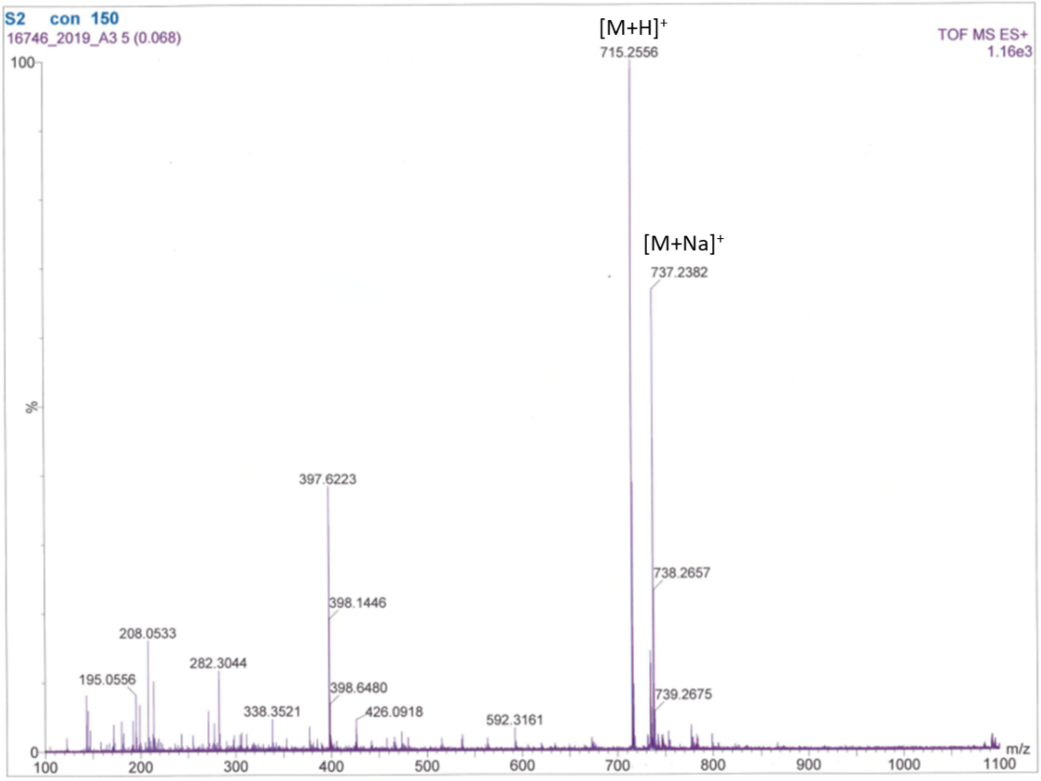


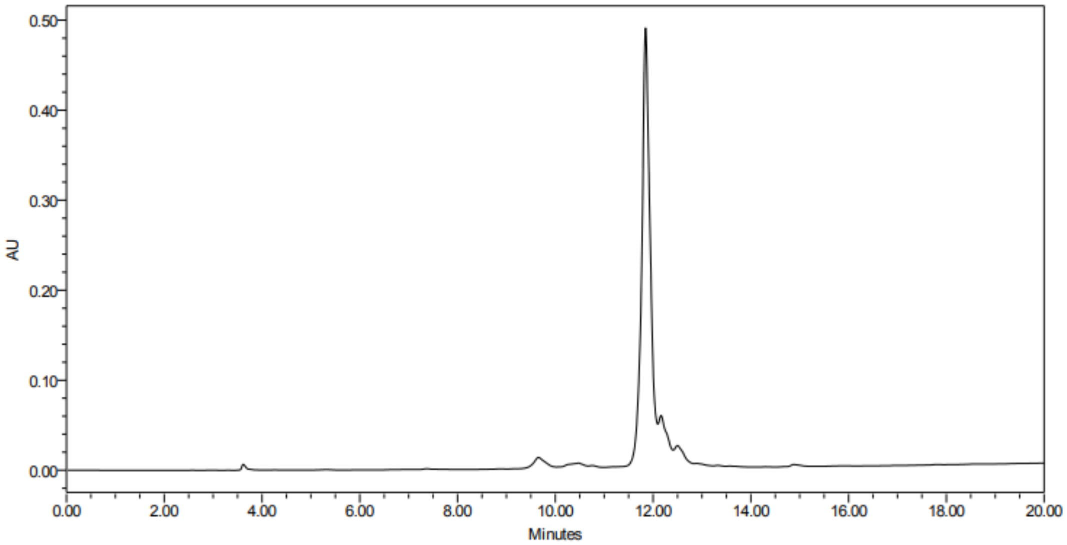

**Ub-ACC**

HRMS for $C_{389}H_{637}N_{107}O_{120}S$

| charge | m/z $_{calcd}$ | m/z $_{found}$ |
|---|---|---|
| +6 | 1461.6221 | 1461.8839 |
| +7 | 1252.9628 | 1253.2618 |
| +8 | 1096.4684 | 1096.7080 |
| +9 | 974.7505 | 974.9593 |
| +10 | 877.3762 | 877.6428 |
| +11 | 797.7063 | 797.8460 |

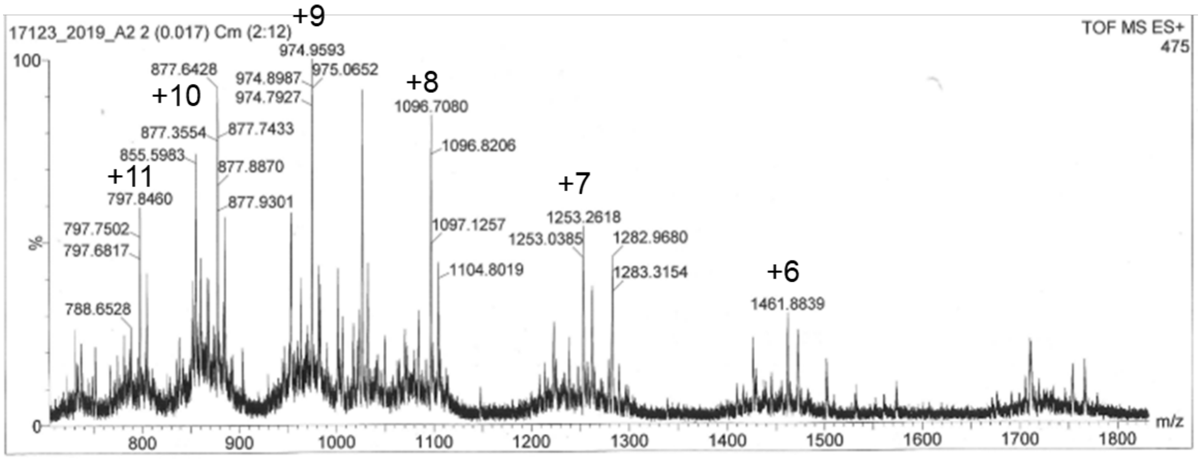


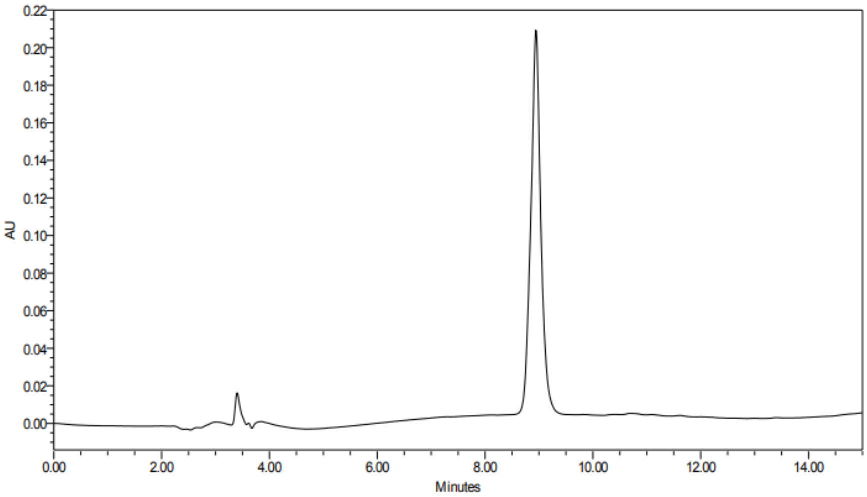

**Ub-Tle-Phg-Gly-Gly-ACC**

HRMS for $C_{391}H_{632}N_{104}O_{120}S$

| charge | m/z $_{calcd}$ | m/z $_{found}$ |
|---|---|---|
| +5 | 1749.1354 | 1749.5309 |
| +6 | 1457.7807 | 1457.9774 |
| +7 | 1249.6702 | 1249.8422 |
| +8 | 1093.5874 | 1093.7565 |
| +9 | 972.1896 | 972.3176 |
| +10 | 875.0713 | 875.1980 |

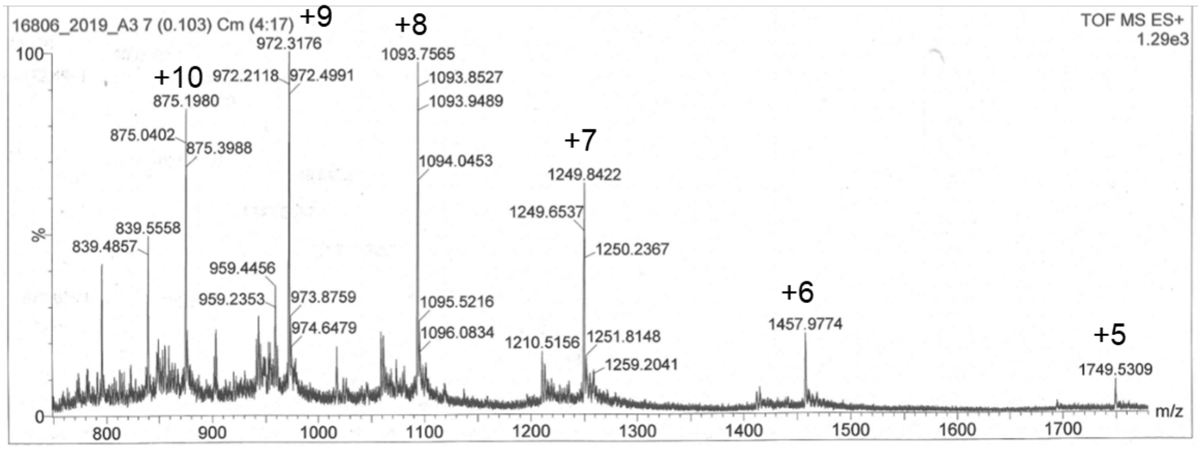


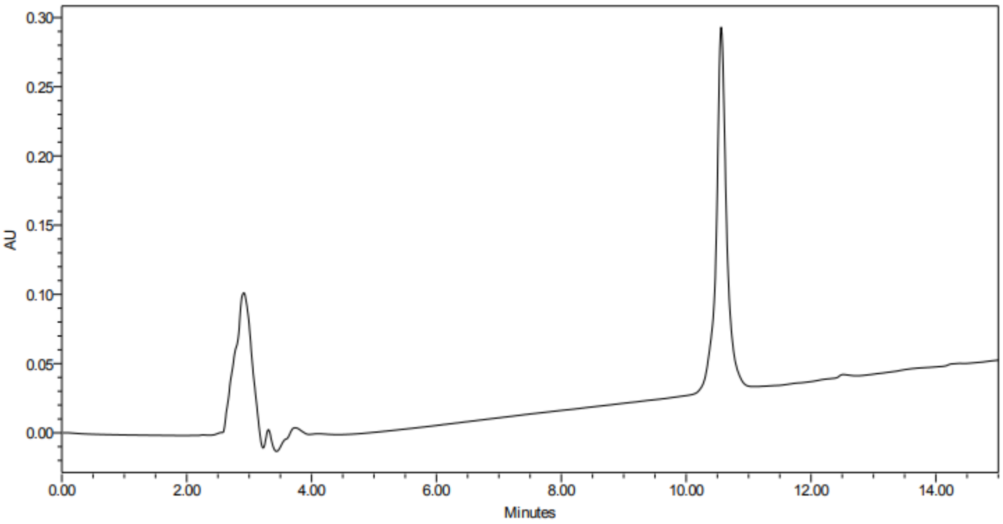

**Ub-Cha-Arg-Abu-Gly-ACC**

HRMS for $C_{394}H_{645}N_{107}O_{120}S$

| charge | m/z calcd | m/z found |
|---|---|---|
| +5 | 1767.3576 | 1767.8867 |
| +6 | 1472.9659 | 1473.3879 |
| +7 | 1262.6861 | 1263.0165 |
| +8 | 1104.9762 | 1105.1364 |
| +9 | 982.3130 | 982.4749 |
| +10 | 884.1824 | 884.3774 |
| +11 | 803.8938 | 804.0812 |
| +12 | 736.9866 | 737.1602 |

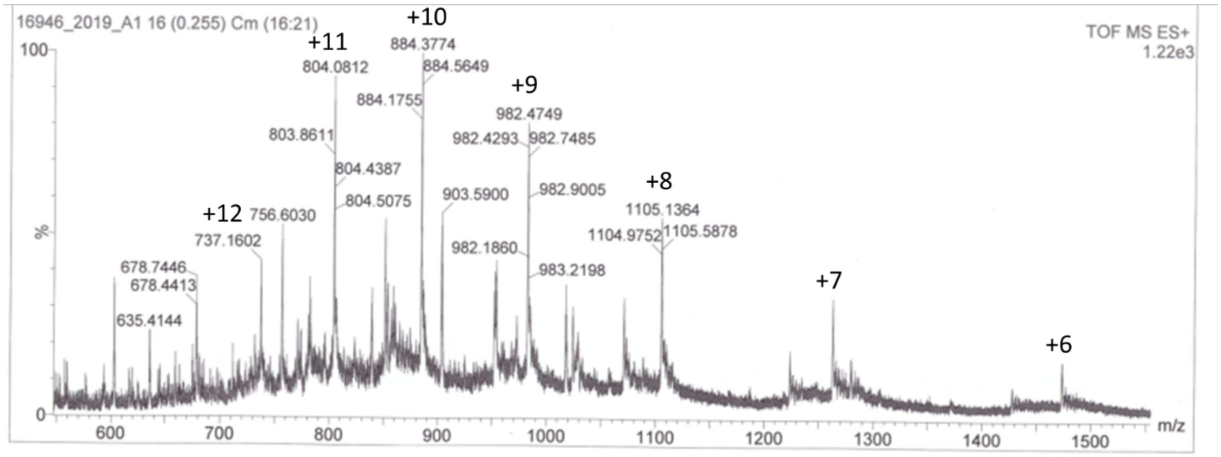


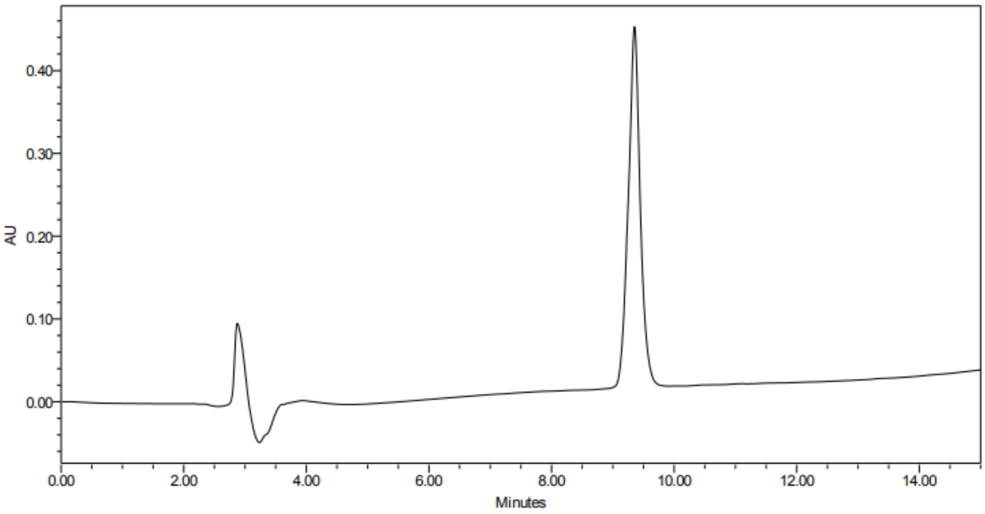

**Ub-DArg-Phe(guan)-Ala-Gly-ACC**

HRMS for $C_{394}H_{640}N_{110}O_{120}S$

| charge | m/z $_{calcd}$ | m/z $_{found}$ |
|---|---|---|
| +4 | 2218.1876 | 2218.8503 |
| +5 | 1774.7516 | 1775.1068 |
| +6 | 1479.1275 | 1479.4045 |
| +7 | 1267.9675 | 1268.1271 |
| +8 | 1109.5975 | 1109.8116 |

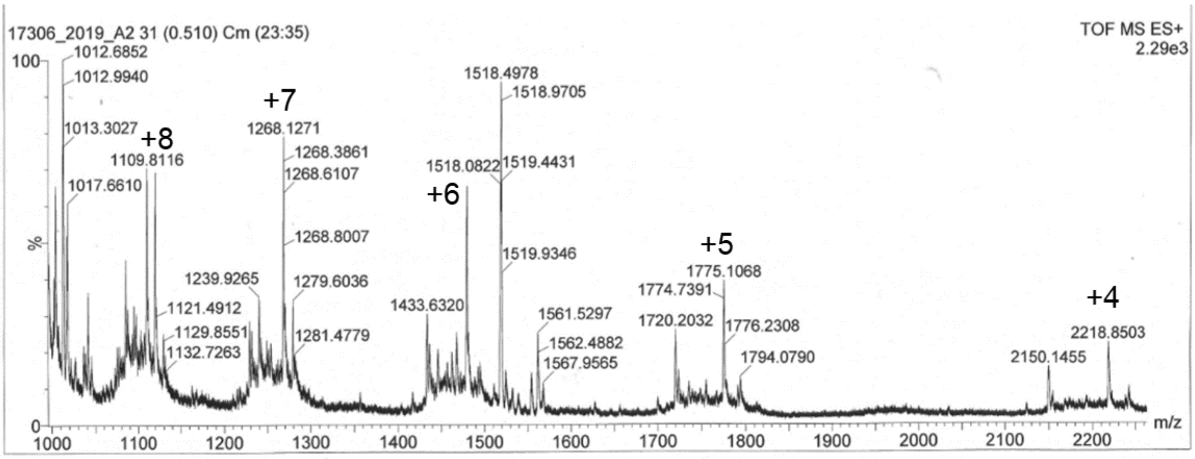


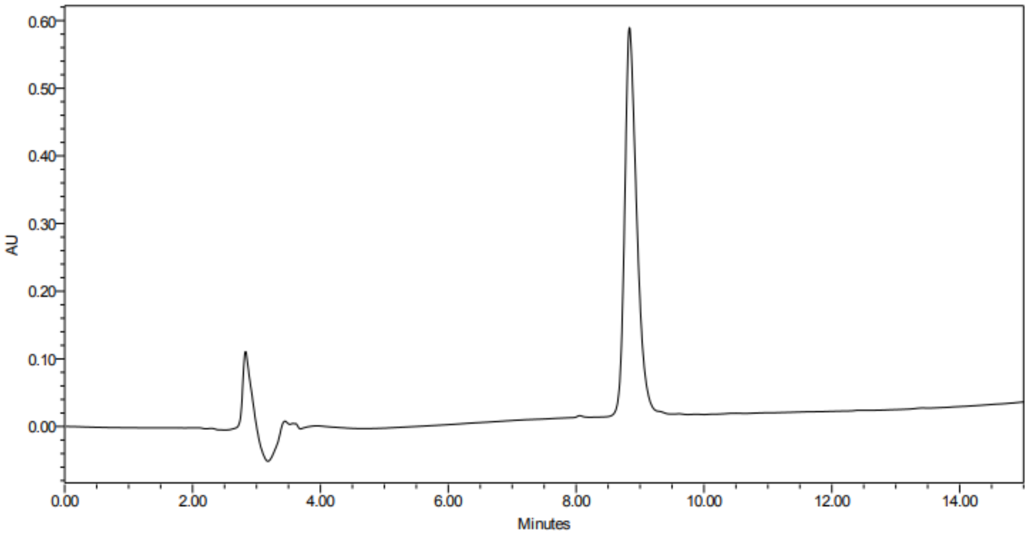

**Biot-6-Ahx-Ub-VME**

HRMS for $C_{397}H_{658}N_{108}O_{121}S_2$

| charge | m/z calcd | m/z found |
|---|---|---|
| +6 | 1491.4778 | 1491.8063 |
| +7 | 1278.5535 | 1278.8268 |
| +8 | 1118.8602 | 1119.2485 |
| +9 | 994.6543 | 994.8820 |
| +10 | 895.2896 | 895.3971 |
| +11 | 813.9912 | 814.0791 |
| +12 | 746.2426 | 746.3508 |

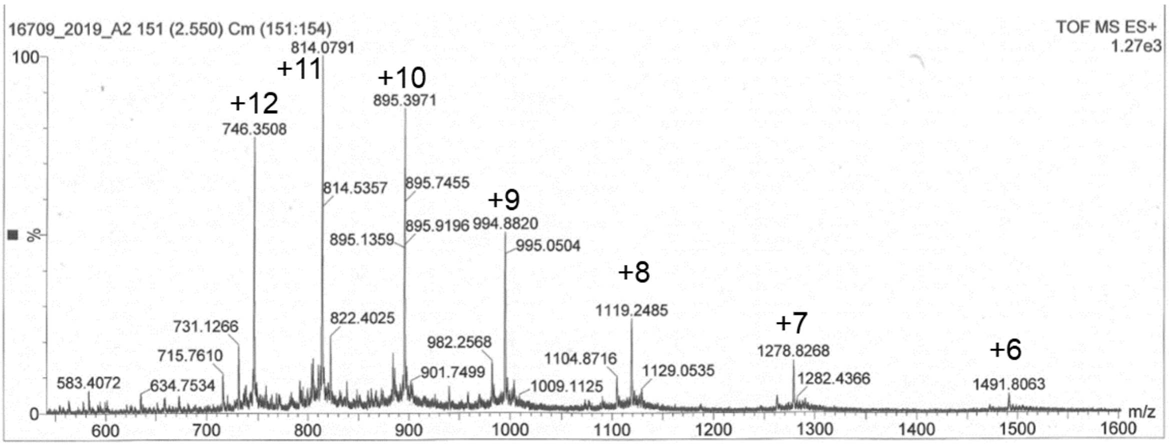


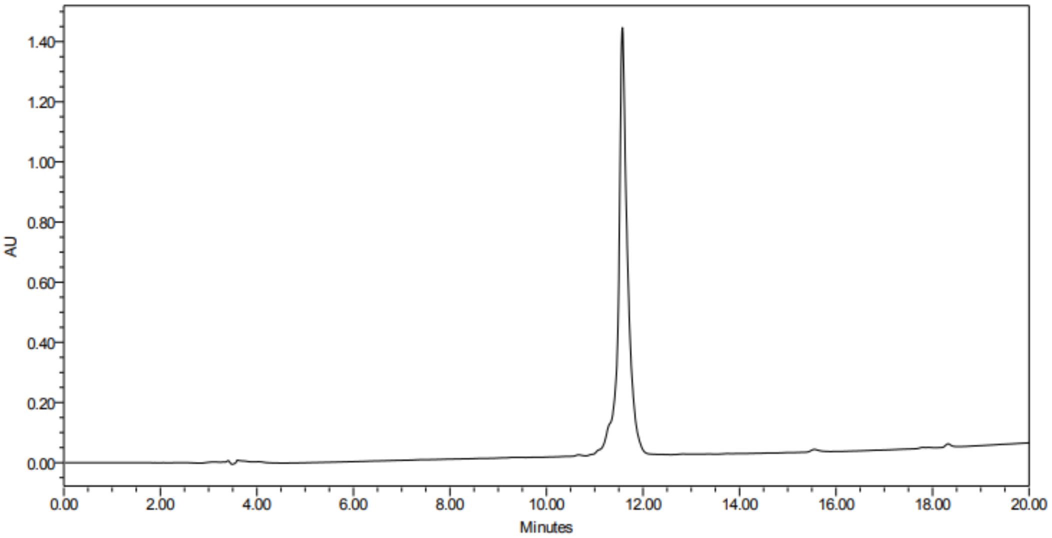

**Biot-6-Ahx-Ub-Tle-Phg-Gly-Gly-VME**

HRMS for $C_{399}H_{653}N_{105}O_{121}S_2$

| charge | m/z calcd | m/z found |
|---|---|---|
| +6 | 1487.6364 | 1487.8646 |
| +7 | 1275.2608 | 1275.4232 |
| +8 | 1115.9791 | 1116.2777 |
| +9 | 992.0934 | 992.2054 |
| +10 | 892.9848 | 893.1783 |
| +11 | 811.8959 | 812.1168 |

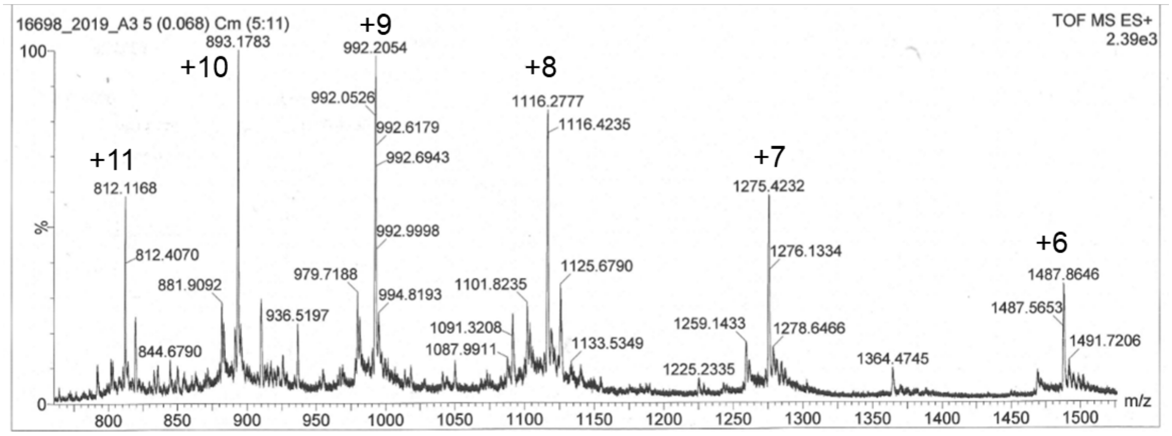


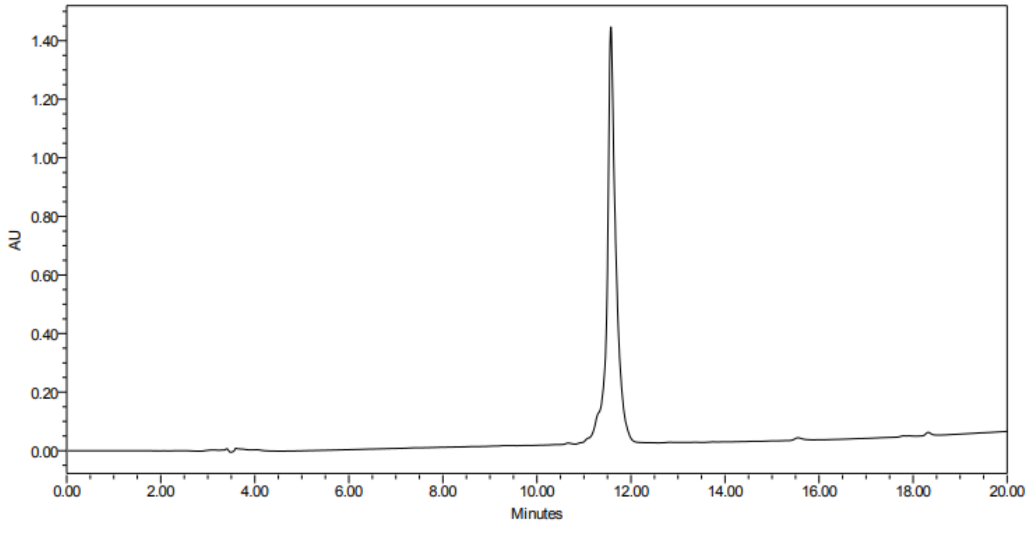

**Biot-6-Ahx-Ub-Cha-Arg-Abu-Gly-VME**

HRMS for $C_{402}H_{666}N_{108}O_{121}S_2$

| charge | m/z calcd | m/z found |
|---|---|---|
| +8 | 1127.3680 | 1127.5012 |
| +9 | 1002.2168 | 1002.3268 |
| +10 | 902.0959 | 902.2721 |
| +11 | 820.1787 | 820.2746 |
| +12 | 751.9144 | 752.0181 |

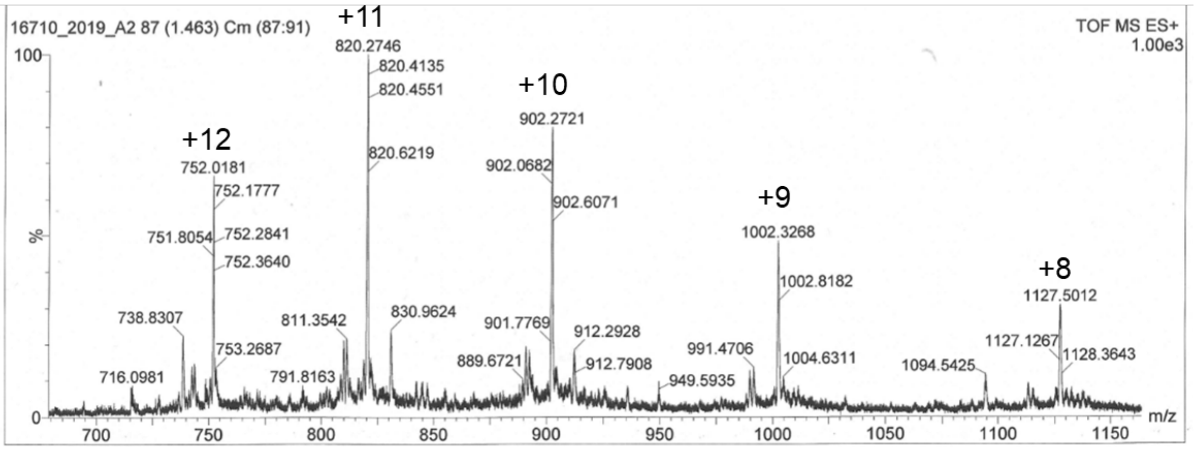


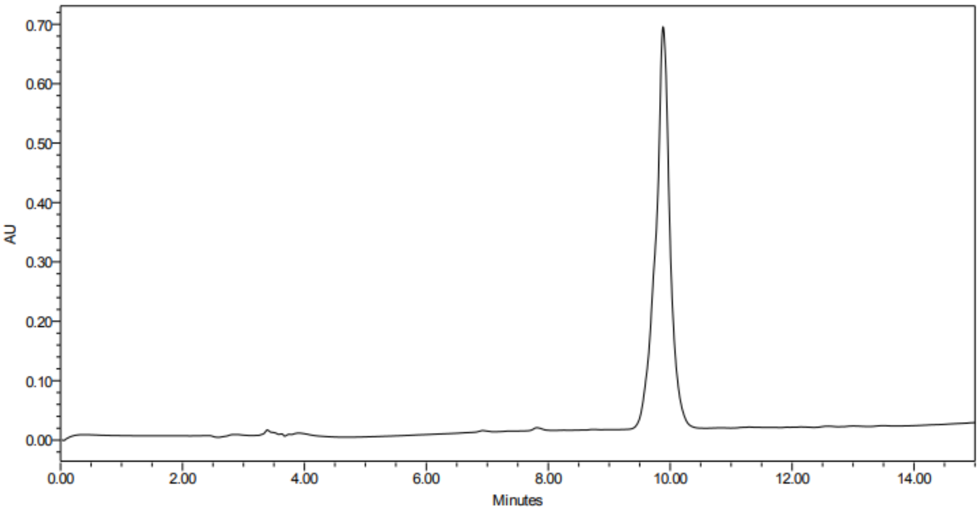

**Biot-6-Ahx-Ub-DArg-Phe(guan)-Ala-Gly-VME**

HRMS for $C_{402}H_{661}N_{111}O_{121}S_2$

| charge | m/z calcd | m/z found |
|---|---|---|
| +8 | 1131.9893 | 1132.2377 |
| +9 | 1006.3246 | 1006.6258 |
| +10 | 905.7929 | 905.9586 |
| +11 | 823.5396 | 823.6242 |
| +12 | 754.9953 | 755.0801 |
| +13 | 696.9962 | 697.1877 |

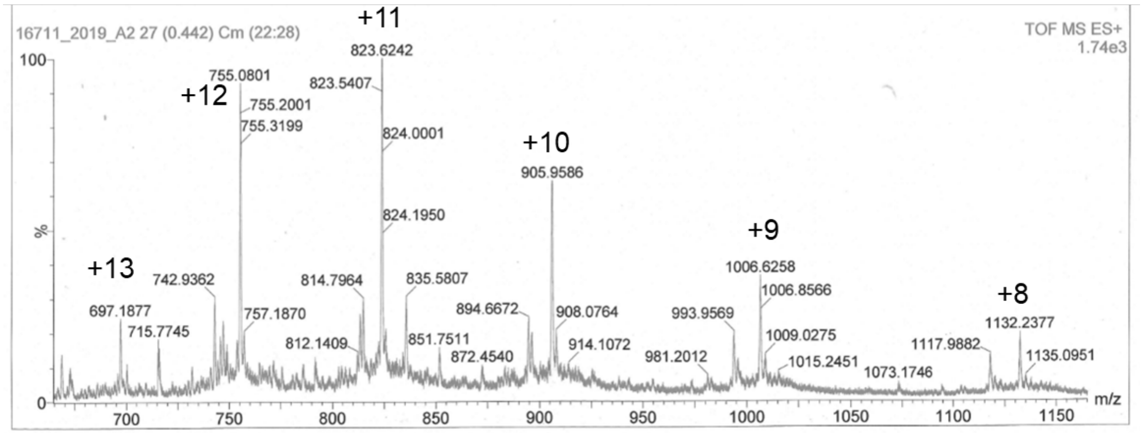


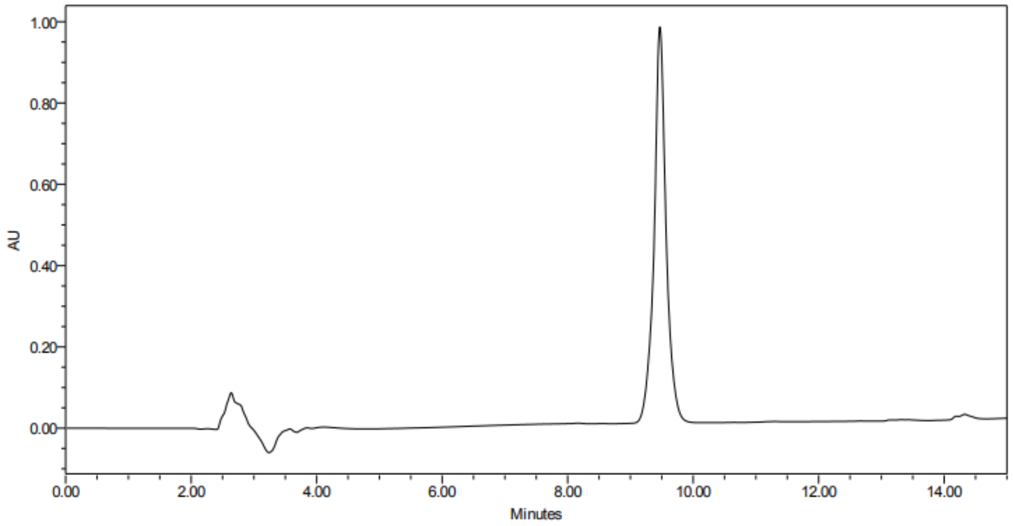
